# Divergent neuronal DNA methylation patterns across human cortical development: Critical periods and a unique role of CpH methylation

**DOI:** 10.1101/428391

**Authors:** AJ Price, L Collado-Torres, NA Ivanov, W Xia, EE Burke, JH Shin, R Tao, L Ma, Y Jia, TM Hyde, JE Kleinman, DR Weinberger, AE Jaffe

## Abstract

We have characterized the landscape of DNA methylation (DNAm) across the first two decades of human neocortical development in NeuN+ neurons using whole-genome bisulfite sequencing and compared them to non-neurons (primarily glia) and prenatal homogenate cortex. We show that DNAm changes more dramatically during the first five years of postnatal life than during the entire remaining period. We further refined global patterns of increasingly divergent neuronal CpG and CpH methylation (mCpG and mCpH) into six developmental trajectories and found that in contrast to genome-wide patterns, neighboring mCpG and mCpH levels within these regions were highly correlated. We then integrated paired RNA-seq data and identified direct regulation of hundreds of transcripts and their splicing events exclusively by mCpH levels, independently from mCpG levels, across this period. We finally explored the relationship between DNAm patterns and development of brain-related phenotypes and found enriched heritability for many phenotypes within DNAm features we identify.

## Introduction

Neurons are unique cells that persist throughout the lifespan, accumulating programmed developmental changes and environmental experience that fine tune neural circuitry in the brain. During development and maturation, neurons undergo precisely coordinated cascades of genetic regulation that combine with experience to shape the cellular output via progressive changes to the epigenome. DNA methylation (DNAm) is an integral facet of the epigenome that plays a role in establishing cell identity and developmental trajectories as well as adapting to experience via regulation of gene expression. Previous large-scale studies of DNAm across human brain development have been limited to homogenate tissue^1^ or have used microarray technologies^2^, creating ambiguity about the extent of cell type-specific developmental DNAm changes and effects on transcript isoforms across the genome^3^.

To better characterize the DNAm landscape across human cortical development, we performed whole-genome bisulfite sequencing (WGBS, see Methods) on homogenate tissue and on a neuron-enriched population isolated from 24 human dorsolateral prefrontal cortex (DLPFC) samples aged 0-23 years using NeuN-based fluorescence-activated nuclear sorting (FANS, **Figure S1A**). To complement these data, we sequenced eight FANS-derived NeuN- postnatal samples and 20 homogenate prenatal cortical samples, for a total of 75 samples after quality control (**Table S1**). We fully characterized the landscape of DNAm at both CpG and non-CpG (CpH) dinucleotides in these samples, allowing for a finer dissection of differential DNAm functional specificity. We also sequenced matched transcriptomes of homogenate cortical samples from these donors and a subset of three nuclear transcriptomes each from NeuN+ and NeuN- samples to assess the functional consequences of epigenomic remodeling (53 total transcriptomes, **Table S2**). By exploring DNAm patterns in neurons across prenatal and postnatal human brain development, we show that the first five years of postnatal life are a critical period in epigenetic plasticity, and we identify developmental shifts in neuronal DNAm in both the CpG and CpH context. We also clarify the relationship between CpG but particularly CpH methylation (mCpG and mCpH, respectively) and gene expression and splicing in neuronal development, and explore the ramifications of these insights for neuropsychiatric disease.

## Results

After data processing, quality control and filtering, we analyzed 18.7 million cytosines in the CpG context at an average coverage of 15x (see methods). Comparable to previous reports^1,4,5^, CpGs were overall highly methylated (71-76% CpGs with ***β***>80%, **Table S3**).

While NeuN antibody labels most mature neuronal subtypes in human cortex, some neurons will not be labeled and will be captured in the NeuN- fraction amidst a diverse array of non-neuronal cell types, including oligodendrocytes, astrocytes, microglia, and epithelial cells. Gene expression differences between fractions confirmed, however, that NeuN+ and NeuN- samples are enriched for neuronal and glial-lineage cells, respectively (**Figure S1B-D**). Therefore in this work we refer to NeuN+ and NeuN- samples as “neurons” and “glia” although we acknowledge that these samples do not perfectly reflect these identities and mask more granular differences between subcellular identities contained within.

Developmental DNAm changes identified in homogenate cortex were strongly confounded by shifting cell type proportions (OR=7.5, p<10^−100^, **Figure S2A**)^2^. While homogenate measurements were positively correlated with developmental changes that occurred in both neuronal and glial cell types (ρ=0.79, p<10^−100^), cell type-specific developmental changes were less consistently observed in homogenate preparations (ρ=-0.26, p<10^−100^, **Figure S2B-D**). Overall, ~40% of cell type-specific developmental DNAm changes could not be detected at all in homogenate cortex (**Figure S2E**), and many of the cell type-specific effects could not be accurately identified in homogenate tissue. These results highlight the importance of measuring DNAm in the appropriate cellular context for improved resolution to detect true developmental changes.

### DNAm as a map of putative functional genomic states

Local CpG methylation (mCpG) patterns are known to distinguish genomic states of DNA and chromatin. For instance, unmethylated regions (UMRs) are associated with promoters, with a subset of longer UMRs (DNAm valleys, DMVs) that overlap developmental genes often encoding transcription factors (TFs)^6,7^; low-methylated regions (LMRs) often signify enhancer sequence^8^; and partially methylated domains (PMDs) are associated with heterochromatin and late replicating DNA^9–11^. To better resolve the developing regulatory landscape in postnatal neurons and glia and in bulk prenatal cortex, we assessed the temporal dynamics of these selected DNAm patterns in the CpG context. Compared to prenatal homogenate cortex and postnatal glial cells, postnatal neurons showed a general accumulation of mCpG, at a rate 50% faster than the other cells. This was evident in the LMR and to a lesser extent the UMR landscape, since fewer and smaller LMRs were identified as neuronal development progressed (**Figure S3A-B**). As expected, UMRs and LMRs were highly enriched for transcription start sites (TSSs) and enhancers in DLPFC chromatin state data from the Roadmap Epigenomics Consortium^12^ (**Figure S3C**). Interestingly, LMRs were similarly enriched in these states in both adult and fetal brain; this correspondence may reflect a shared regulatory landscape established early in development.

While PMDs are a common feature of most cell types, they have not been conclusively identified in neurons. Here we identified a range of 245 to 404 PMDs per neuronal sample (**Figure S4A**). PMDs were especially enriched for heterochromatin and, interestingly, enhancers in our postnatal neuronal samples (**Figure S4B**). 65.4% of PMD base pairs were also identified as PMD in an independent WGBS dataset of NeuN-sorted human neurons (**Figure S4C**). 40.3-61.0% of PMD bases per neuronal sample were identified as common PMD sequence, and 9.3-15.0% bases were additionally identified as PMD in at least one sample in a recent study profiling PMDs in multiple cell types and tissues^13^ (**Figure S4D**). These data suggest that although the neuronal genome was overall highly methylated, a small but consistent portion displayed the characteristics of PMDs.

We further identified significant neuronal DMV changes through accumulation of mCpG that revealed regulators of cell identity and development and their temporal windows of expression change. Compared to bulk prenatal cortex, postnatal neurons and glia showed marked reduction in the size of DMVs (**Figure S5A**). Methylation shifts within DMVs led to inclusion and exclusion of transcription factor genes in an age-dependent manner, and on average transcription factor genes were higher expressed in the age group in which the gene was escaping the DMV state by accumulating DNAm (**Figure S5B-C**). These results underscore the substantial DNAm landscape alterations that neurons and glia undergo during development in defined mCpG patterns, including previously unobserved PMDs.

### Developmental shifts in neuronal mCpG highlight synaptic remodeling during first five years of postnatal life

We next quantified more localized changing mCpG levels by exploiting the correlation between neighboring mCpG levels to identify genomic regions with changing mCpG. We identified 11,179 differentially methylated regions (DMRs, FWER<5%, see Methods) in the CpG context between cell types (covering 31.1 Mb) that replicated in independent WGBS data^1^ (98.4% concordant, ρ=0.925, **Figure S6A**). Many of these DMRs overlapped genes involved in neuronal or glial-specific processes (**Figure S6B**). We found fewer DMRs for developmental mCpG changes compared with cell type differences, the majority being within rather than across cell types (2,178 versus 129 DMRs, at ~5% change in DNAm per decade of life, FWER<5%). Among the 2,178 cell type-specific developmental DMRs (cdDMRs, 3 Mb, **Table S4**), neuronal mCpG patterns seemed to diverge from an immature landscape shared by glia and prenatal cortex (**Figure 1A**), with the largest changes occuring in the first five years of life. Indeed, the magnitude of DNAm changes in neurons and glia in samples five years and younger was double that of older samples (**Figure S6C-D**). These results provide epigenetic correlates to the known developmental processes occuring in the cortex in the first five postnatal years, including prolific synaptogenesis and gliogenesis.

**Figure 1:**
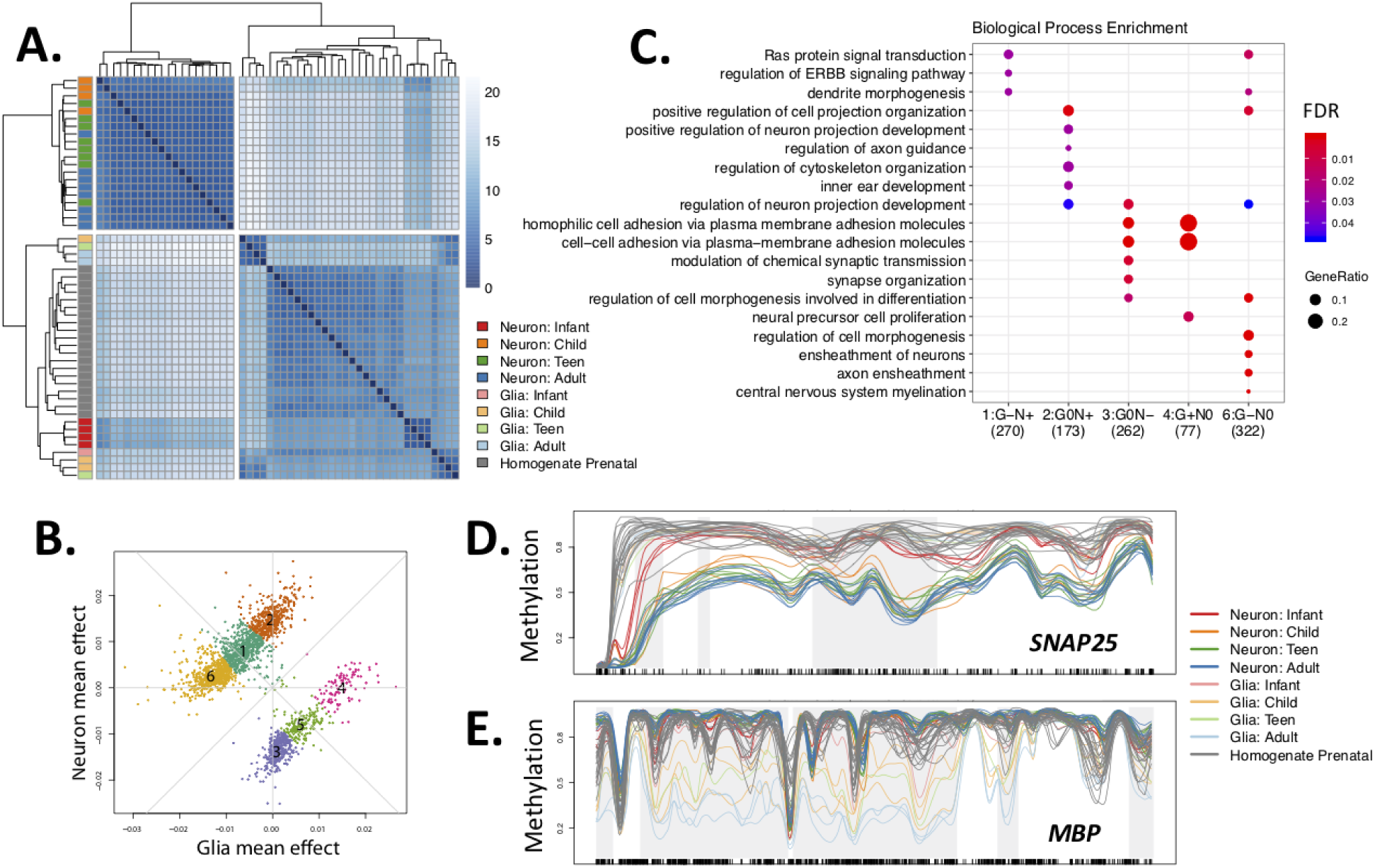
Regional cell type-specific developmental mCpG trajectories. **(A)** Euclidean distances between samples within cdDMRs shows that older neuronal samples cluster separately from infant neuronal samples, glia regardless of age, and bulk prenatal cortex. **(B)** Decomposing cdDMRs patterns into 6 clusters using *k-means* based on glia and neuron mean mCpG changes per year of life. **(C)** The top five most enriched gene ontology terms for each of the six groups in **(B)** highlights diverse biological processes amongst the groups. No terms were enriched for Group 5. **(D)** Example Group 3 cdDMR within *SNAP25*. **(E)** Example Group 6 cdDMR within *MBP*. Gray shading indicates the boundaries of the cdDMR, and black tick marks on the x-axis indicate the position of CpGs. Key: Neuron (NeuN+); Glia (NeuN-); Infant (0-1 years); Child (1-10 years), Teen (11-17 years) and Adult (18+ years).

We further parsed these cdDMRs using k-means clustering to partition the cdDMRs into six groups with unique DNAm characteristics (**Figure 1B**). 71.1% of cdDMRs were in groups characterized by increasing neuronal and/or decreasing glial DNAm over postnatal development (**Figure 1B;** Groups 1, 2 & 6). A varying proportion of each cdDMR group corresponded to sequence differentially methylated by neuronal subtype from publicly available data^14^ depending on the trajectory of neuronal methylation patterns in the group, suggesting that assorted neuronal subclasses contribute to these developmental patterns (**Figure S7A-B**). Gene ontology enrichment in the six groups suggested that these groups are associated with a continuum of biological roles, many relating to functions specific to the cell type with decreasing methylation (**Figure 1C**). For example, **Figure 1D** shows a Group 3 cdDMR within *SNAP25*, a presynaptic neuronal gene, in which neurons uniquely and progressively lost DNAm over development. This pattern suggests increased repression of a neuronal fate in maturing glia not mirrored in neurons over postnatal development in Group 3 cdDMRs. Likewise, the opposite pattern was observed in a Group 6 cdDMR within *MBP*, an oligodendrocyte gene encoding a component of the myelin sheath, in which glia but not neurons progressively lost DNAm (**Figure 1E**).

We lastly compared these cdDMR groups to a list of putative enhancers active in human brain development curated by evolutionary age^15^ and found strong enrichment for these sequences across all six groups (**Figure S8A**). Human accelerated regions, or conserved sequences that have experienced rapid mutation in the human lineage^16^, were also enriched for dynamic DNAm remodeling (**Figure S8B**), suggesting that our CpG-based cdDMRs may be enriched for sequences related to higher cognitive functions associated with the human DLPFC.

Overlapping cdDMRs with the mCpG features identified above provided additional insight to the potential functional genomic states underlying these regions. For instance, cdDMRs scarcely overlapped heterochromatic PMDs; cdDMRs losing neuronal mCpG were positively correlated with increasing LMR overlap, potentially reflecting enhancer element activation during cortical maturation in these groups (Groups 3 & 5 cdDMRs; both with t>3.8, ρ>0.63, and FDR<2.7e-03). Curiously, a high proportion of cdDMRs gaining DNAm in glia but not in neurons (Group 4 cdDMRs) overlapped DMVs early in development in glia but steadily lost DMV status over time (t=-4.3, ρ=-0.87, FDR=1.3e-02, **Figure S9A**). Assessing chromatin state from the homogenate Roadmap Epigenomics brain maps, in contrast, lacked the resolution to provide this nuance: all six cdDMR groups were similarly enriched for transcriptional (particularly TSS-flanking) and enhancer chromatin states, and depleted for heterochromatin and quiescent states (**Figure S9B**). These results confirm the role of dynamic DNAm in helping establish epigenomic states that guide cell lineage differentiation and emphasize the utility of creating genome-wide DNAm maps to better parse the functional diversity of cell type-specific developmental DNAm remodeling in the human cortex, a process that is particularly critical during the first five years of postnatal development.

### Abundant neuronal CpH methylation is highly correlated with neighboring CpG methylation

Unlike in most other somatic tissues and cell types, mCpH is an abundant, conserved feature of the neuronal epigenome^1,4^. We therefore analyzed 58.1 million cytosines in CpH contexts (H=A,T, or C) that had evidence of methylation across the samples (coverage ≥5, at least 5 samples with ***β***>0, see Methods). As shown previously^1^, mCpH sites were predominantly lowly methylated (92-99% CpHs with ***β***<20%, **Table S3**). While mCpH was distributed throughout the genome (**Figure S10A**), it was greater in neurons than glia (98.9% of 7,682,075 differentially methylated CpHs were in neurons, at FDR<5%) and mostly accumulated across postnatal development (99.3% of 3,194,618 CpHs, at FDR<5%; **Table S5**). Most mCpH accumulated primarily in either the CAG or CAC context over the first five years of postnatal life—similarly to mCpG—followed by a tapered global increase into adulthood (**Figure S10B**).

While the majority of mCpH in embryonic stem cells (ESCs) occurs in the CAG context, previous work has shown that ESCs undergo loss of mCAG during neuronal differentiation followed by preferential accumulation of mCAC^17^. Here we further refined these patterns and found a cell type-specific relationship with trinucleotide context: overall, total mCAG increased 40% faster than mCAC in neurons, while in glia, mCAG accumulated 50% slower than mCAC (**Figure S10C**). Taking into account relative genome-wide proportions of CAG and CAC though, neuronal mCAG accumulated 30% slower than mCAC (**Figure 2A**). mCH that was greater in glia than neurons, or in younger than older neurons, was more likely to be in the CAG than CAC context (OR>4.13, p<2.2e-16). Interestingly, the 3,286 and 1,744 genes that contained significantly increasing and decreasing mCAC versus mCAG over development, respectively, were associated with different biological processes related to neuronal function and activity, particularly involving the synapse (**Figure S10D**). These results reinforce that trinucleotide context may be regulated by distinct mechanisms playing non-redundant biological roles in human brain development.

**Figure 2:**
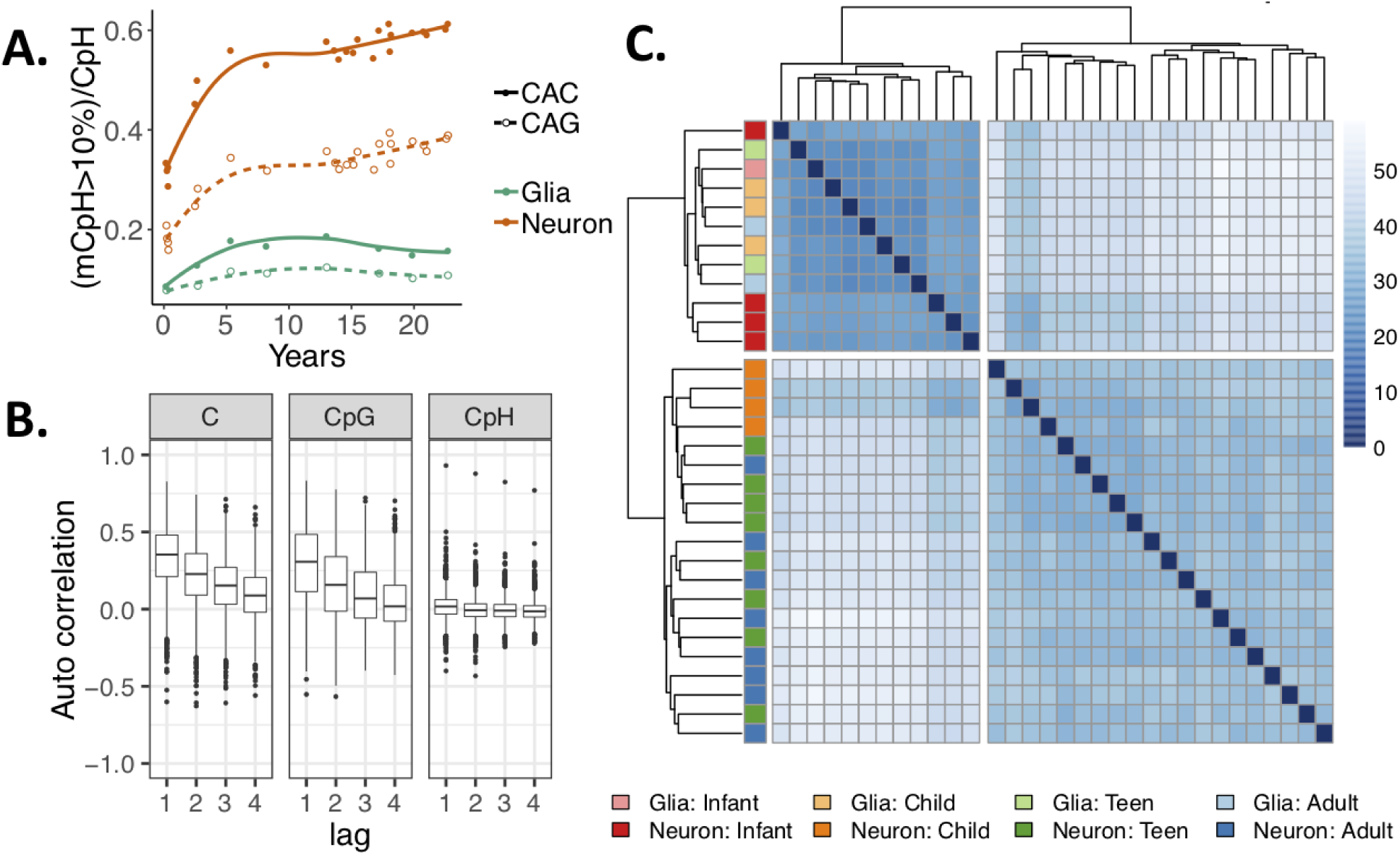
CpH methylation patterns across brain development. **(A)** The proportion of CAC and CAG sites that are greater than 10% methylated in neurons and non-neurons (glia) across brain development. **(B)** Autocorrelation levels for different cytosine contexts in neurons. Autocorrelation levels were similar for mCpG and all cytosines, with uncorrelated levels in the CpH context. **(C)** Euclidean distances between samples based on mCpH within cdDMRs again cluster infant neurons dark red) with glia of all ages (light colors) rather than with older neurons.

We next examined the relationship between neighboring mC levels by measuring autocorrelation, or how correlated the methylation level of a cytosine is with that of cytosines progressively further away. Unlike in the CpG context, where neighboring mCpG levels were highly correlated as previously described^18^, neighboring CpH DNAm levels across the genome were not autocorrelated. Within the cdDMRs, however, while mCpH levels separately remained uncorrelated, together all methylated cytosines (i.e., mCpH+mCpG) showed similar autocorrelation as mCpG levels alone (**Figure 2B**). This was especially surprising given that there were about two times as many CpHs than CpGs within these regions and that the CpG and CpH were relatively interspersed, suggesting potential functional convergence in the developmentally regulated patterns identified by mCpG in these regions. Indeed, unsupervised hierarchical clustering of CpH within the cdDMRs showed infant neuronal mCpH levels were even more similar to glia compared to older neurons than mCpG (**Figure 2C**). Examining the mean mCpH compared to mCpG within the k-means cdDMR clusters showed that the groups gaining mCpG were the most correlated with mCpH trajectories within the cdDMRs (ρ**=**0.97, t=17.6, p=2.0e-14), and that although mCpG (unlike mCpH) is present at high levels prenatally, both mCpG and mCpH accumulate at similar rates over postnatal development in these groups, once again especially in the first five years of postnatal life where the majority of the methylation change takes place (t=-0.091, p=0.94; **Figure S11**). These results emphasize the potential regulatory importance of cdDMRs and putative functional agreement between both contexts of DNAm in these regions.

### mCpG and mCpH levels influence transcript isoform use

Previous studies show that both mCpG and mCpH in gene bodies but particularly in the promoter and first 2 kb of the gene are negatively associated with gene expression, and that genic mCpH is the most discriminating predictor of gene expression^1,5^. To anchor our DNAm patterns in transcriptional activity, we compared our WGBS data with NeuN-sorted nuclear RNA-seq data (see Methods). We took the average DNAm levels across six groups—infant (ages 0-1), child (ages 1-10), and teen (age 10+) within both cell types (neuronal and glial)—and calculated associations between DNAm and expression. Gene expression was negatively correlated with mCpG regardless of age and cell type in both promoter sequence and gene bodies (−0.42<ρ<-0.22, p~0; 57,332 genes, p <10^−100^; **Figure S12A**). Interestingly, mCpG in exons was significantly but weakly positively correlated with exon expression in infancy, particularly in glial samples (p=0.094, p <10^−100^), which may relate to the previously-identified positive relationship between mCpG and expression and higher methylation in exons than introns^9^.

Across promoters, gene bodies, and exons, only neurons showed a negative correlation between gene expression and mCpH, a relationship that became stronger over development (**Figure S12B**). This pattern was consistent with the preferential accumulation of mCpH in neurons as the brain matures. mCAC and mCAG showed similar patterns of increasingly strong negative correlation preferentially in neurons between methylation and expression across these features (**Figure S12C-D**). Both mCpG and mCpH surrounding exon-exon splice junctions were weakly negatively correlated with expression of the junction in neurons (**Figure S12E**).

Because mCpG has previously been associated with alternative splicing^19^ and mCpH is 15-20% greater in exons than in introns^9^, we hypothesized that accumulating mCpH may contribute to the diversity of alternative splicing characteristic of the brain particularly during development. Leveraging our single-base resolution data, we were able to identify genome-wide functional correlates of mCpH, independent of nearby mCpG, by associating DNAm with nearby expression in the same cortical samples. Specifically, we tested whether methylation levels directly associated with gene or exon expression levels as well as the “percent spliced in” (PSI) of alternative splicing events using the 22 neuronal samples with matching homogenate polyA+ RNA-seq data (see Methods). We found 40,940 CpG and 40,303 CpH associations that explain changes in these three expression summarizations at FDR<5% with a genome-wide p<5×10^−4^. We further identified 220,622 marginal (p<0.01) CpG associations with expression within 1 kb around the associated CpH. While an independent association of mCpH at the gene and PSI summarizations was rare, there were substantially more exons exclusively regulated by local mCpH, largely in the CHH context, in developing postnatal neurons (**Figure 3A**). Three examples of methylation-associated isoform changes are shown in **Figure 3B**.

**Figure 3:**
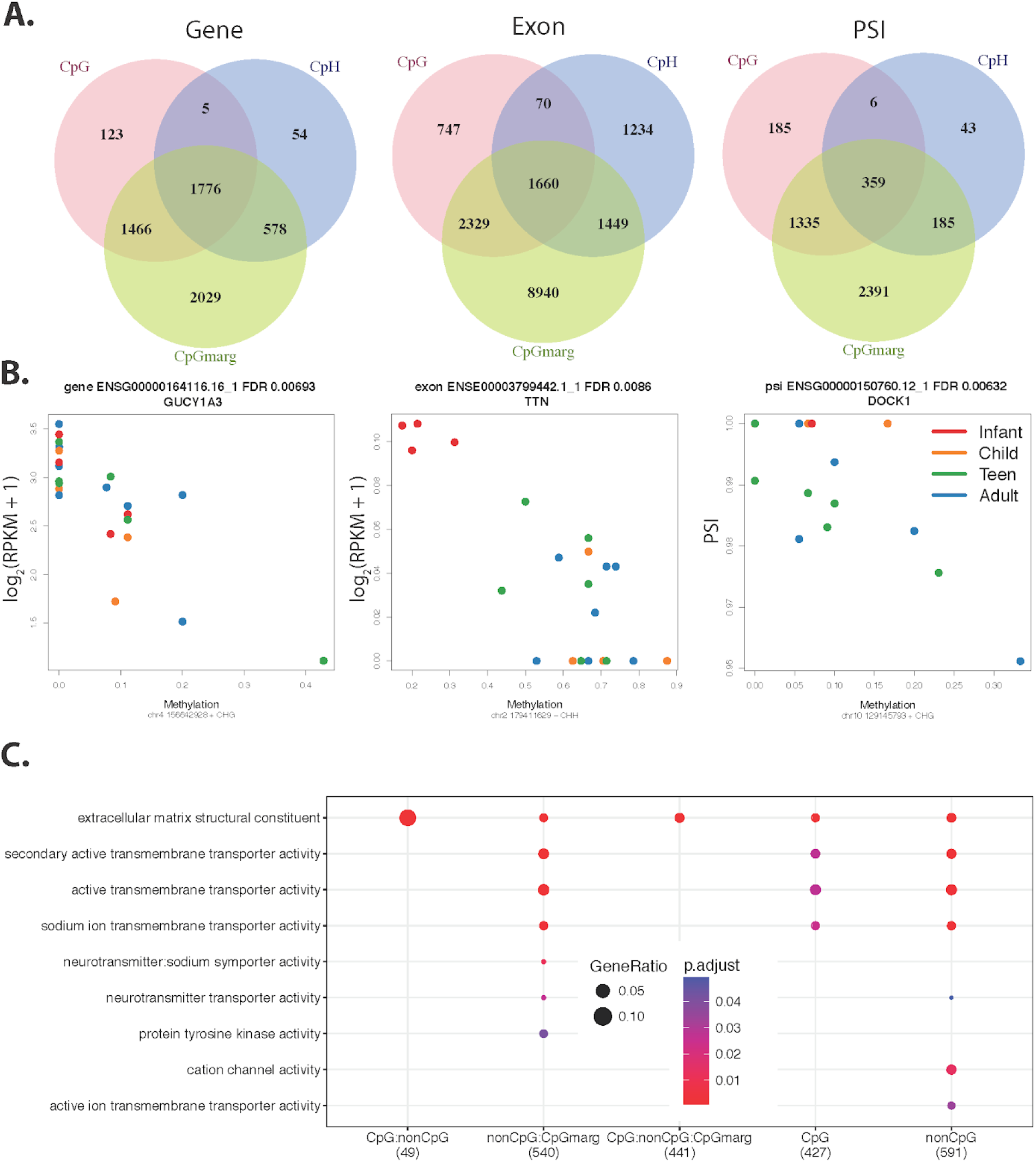
Methylation associations with expression. **(A)** Venn diagrams of the methylation associations by unique feature for the gene, exon and PSI. The sets are determined by if the association is FDR<5% genome-wide for CpG and CpH or if it is a CpG marginally significant within +/- 1 kb window of a CpH association. **(B)** Example associations between methylation and expression at the gene level colored by age: red - infant, orange - child, green - teen, blue - adult. *GUCY1A3* contains one of the top CpH differentially expressed between neurons and glia. Expression of an exon of *TTN*, an autism-associated gene, is negatively associated with mCpH. *DOCK1* PSI of an alternative end site is negatively associated with mCpH. **(C)** Enriched molecular function ontology terms for methylation-associated exons by the venn diagram groups from **(A)**.

Regardless of context specificity, these expression-associated cytosines were depleted in gene promoters and instead enriched in gene bodies and flanking regions (**Table 1**, see Methods). Both contexts were enriched for the high-GC 3’ and 5’ canonical splice site sequences (FDR<1.1e-04), although the associated cytosine could be either inside or outside the corresponding expression feature. Only 3.5-13.7% of expression-associated cytosines overlapped DMR sequence after stratifying by expressed feature and dinucleotide context, indicating that these associations may arise from a more individualized mC effect than the DMRs.

**Table 1:**
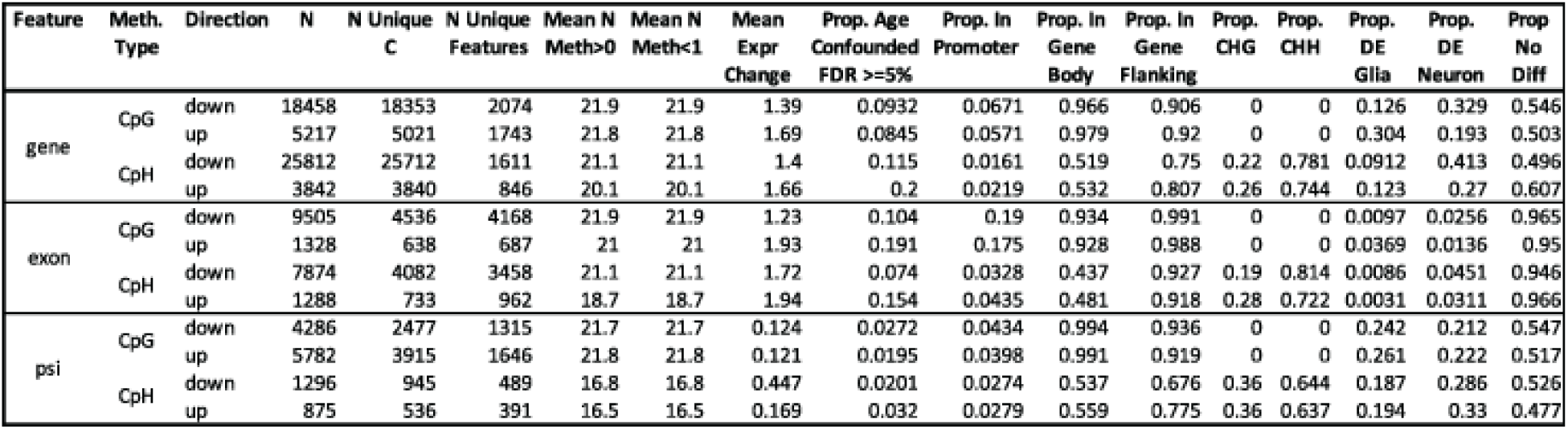
Summary of methylation-expression associations. Mean values are shown for the number of samples with ***β***>0, ***β***<1, and expression change variables (either ΔPSI or Δlog2(RPKM + 1) for genes and exons). Columns from “Age Confounded” are proportions. See Table S10 for a full description of the variables.

Although the majority of these DNAm-expression associations were independent of development despite being identified in developing neurons, the mCpH changes at these sites were independently associated with age and expression. The genes including PSI events regulated by local DNAm levels in both CpG and CpH contexts were consistently enriched for neuronal components (**Figure S13**), while genes containing methylation-associated alternative exons were enriched for synaptic signaling and neurotransmitter transport (**Figure 3C** and **Figure S14**), suggesting that we are detecting true neuronal mC-expression associations despite measuring splicing in homogenate RNA-seq. Many of these genes were also differentially expressed between neuronal and glial nuclear RNA (FDR<0.05, **Table 1**). Most, but not all, expression-associated cytosines at the gene- and exon-level showed significant decreases in expression as methylation levels increased.

The associations between these putatively regulatory cytosines and nearby expression levels can be explored in a webtool (https://jhubiostatistics.shinyapps.io/wgbsExprs/). Results can be interactively summarized such as in **Table 1** for user-selected subsets and visualized as in **Figure 3B** or via the UCSC genome browser (**Figure S15**). By integrating neuronal mCpG and mCpH levels with accompanying RNA-seq data in the same brains, we have identified, for the first time, direct regulation of hundreds of transcripts and their splicing events exclusively by mCpH, independent of mCpG levels, across the first two decades of human cortical development.

### DNAm patterns shed light on the active cell type and timing of neuropsychiatric phenotype development

Previous work has attributed a high proportion of neuropsychiatric trait heritability to neuron-specific DNA methylation patterns^20^. Given the role of dynamic DNAm in marking DNA sequence function over development, we examined the relationship between our methylation features with heritability for 30 human behavioral-cognitive traits, psychiatric and neurological disorders, and non-brain-related traits^21^ (**Table S6**), hypothesizing that DNAm patterns may illuminate not only the active cell type but potential critical timeframes for genomic activity in these complex phenotypes. We used stratified linkage disequilibrium score regression (LDSC)^22^ to estimate the proportion of heritability measured in GWAS summary statistics for each phenotype that could be attributed to each of 16 genomic features, including 10 sets of DMRs, LMRs identified in the prenatal, glial or neuronal methylome, human brain regulatory sequence annotated by chromHMM or the LDSC package, or non-differential CpG clusters (**Figure 4A**). In agreement with previous findings^20,23^, human brain annotated regulatory sequence was broadly enriched for heritability of brain-specific traits (14 of 26 brain-associated phenotypes enriched in chromHMM or CNS (LDSC) regions at FDR≤0.05), as were neuronal features (10 of 26 brain phenotypes enriched in neuronal hypomethylated regions, FDR≤0.05; **Table S7**). Significantly differentially hypomethylated neuronal regions had on average 1.85 times higher enrichment scores than non-differential neuronal LMRs, meaning the DMRs explained 1.85x more heritability over regions containing a similar number of SNPs than the LMRs. Interestingly, body mass index (BMI) heritability was enriched in general brain regulatory sequence and hypomethylated neuronal DMRs (FDR≤0.05), consistent with previous evidence linking this metabolic phenotype to regulatory sequence active in cells of the human central nervous system^22^.

**Figure 4:**
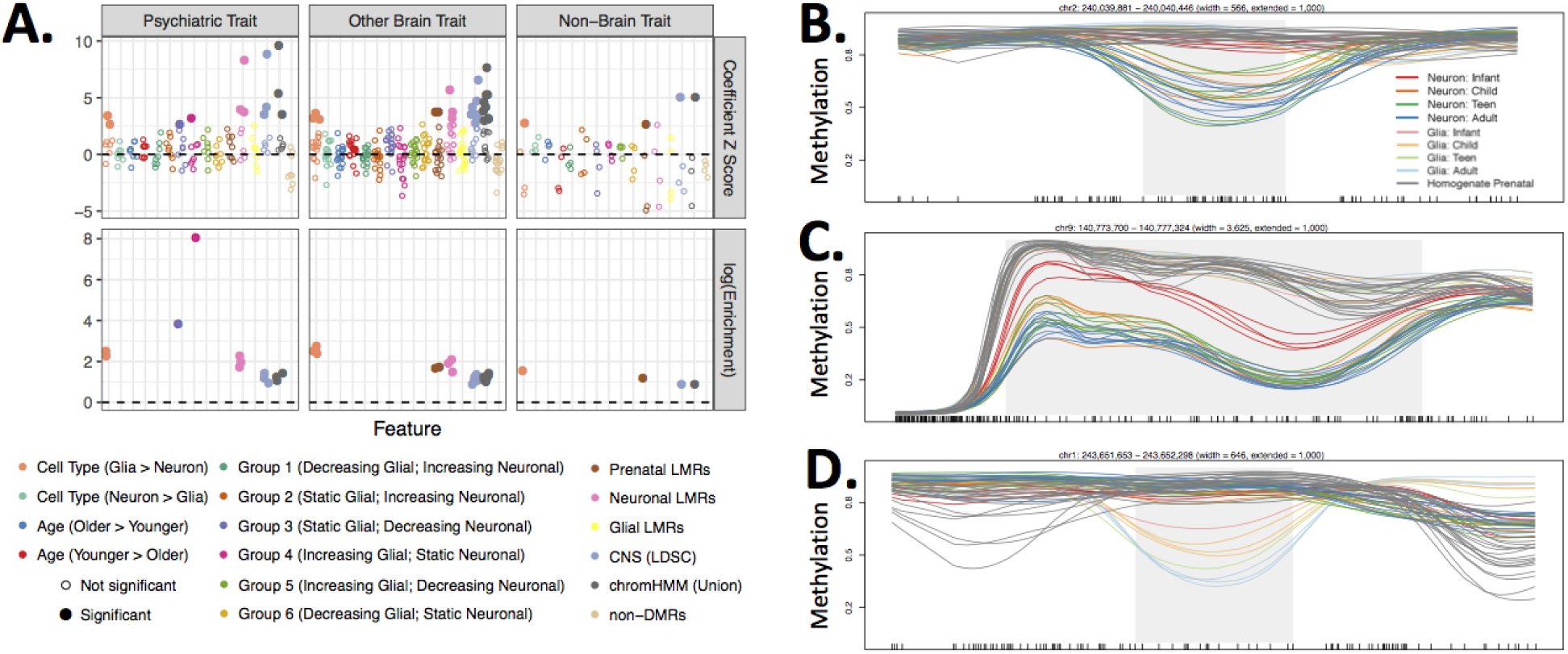
DNAm patterns and brain trait heritability. **(A)** Results assessing enrichment for heritability of 30 phenotypes within 16 groups of DNAm features using Stratified Linkage Disequilibrium Score Regression (SLDSR). Each dot represents results for a single phenotype:DNAm feature pair. The color indicates the DNAm feature, and the phenotypes are stratified by column into psychiatric phenotypes, other brain-related phenotypes (ie, neurological or behavioral-cognitive), or non-brain-related traits. The upper row shows the coefficient Z score for each tested phenotype:DNAm pair, or the amount of additional heritability explained by the DNAm feature over 53 baseline features in the SLDSR model. The lower row shows the enrichment score, or the proportion of heritability attributed to the feature divided by the proportion of SNPs in the feature. For clarity enrichment scores of only the significant feature-trait combination are depicted. Filled in circles indicate significantly enriched heritability for a phenotype in a feature (coefficient p-value corrected using Holms method≤0.05). **(B)** A cdDMR overlapping *HDAC4*, a gene associated with autism spectrum disorder (ASD), shows the Group 3 pattern of decreasing neuronal and static glial DNAm. **(C)** A cdDMR overlapping *CACNA1B*, a gene associated with ASD, shows the Group 5 pattern of decreasing neuronal and increasing glial DNAm. **(D)** A cdDMR overlapping *AKT3*, a gene associated with schizophrenia, shows the Group 6 pattern of decreasing glial and static neuronal DNAm. Gray shading indicates the boundaries of the cdDMR, and black tick marks on the x-axis indicate the position of CpGs.

In terms of developmental DNAm patterns, heritability of BMI, IQ, neuroticism, and major depressive disorder was enriched in both postnatal neuronal and prenatal LMRs, suggesting early action of genetic influence on the development of these phenotypes (FDR≤0.05). Few developmental differential groups captured a significant proportion of heritability for the 30 traits tested, perhaps because of their small size compared to the cell type-specific or non-differential groups (cdDMRs covered ~31, 240 and 838 times less sequence and included ~32, 273 and 904 times fewer SNPs than cell type DMRs, LMRs or general brain features, respectively). However, despite covering only 607 kilobases, group 3 cdDMRs (ie, static glial and decreasing neuronal DNAm) were significantly enriched for heritability of schizophrenia (coefficient z-score=2.74, FDR=0.039). Group 4 cdDMRs (111.7 kilobases; increasing glial, static neuronal DNAm) were also enriched for heritability of PTSD (coefficient z-score=3.21, FDR=0.01).

Given the enrichment for psychiatric disease heritability measured in common SNPs in these cdDMR groups, we then expanded our analysis to include seven curated gene sets containing *de novo* and rare inherited variation--including rare copy number variants (CNVs) and syndromic variants--associated with psychiatric, neurodevelopmental and neurodegenerative disorders^24,25^. We again found significant enrichment of hypomethylated neuronal DMRs in genes implicated in psychiatric and neurodevelopmental disorders (i.e., schizophrenia, autism spectrum disorder (ASD), syndromal neurodevelopmental disorders, and intellectual disability; all with OR>2.04 and FDR<1.9e-02; **Table S8**). In this analysis, we also found enrichment for hypermethylated neuronal DMRs in ASD genes from the SFARI Gene database and schizophrenia genes containing *de novo* mutations (both with OR>1.92 and FDR<5.0e-03). These results confirmed a prominent role of neuronal functioning in most of the neurodevelopmental disorders using an orthogonal measurement approach as done previously^23^. Over postnatal development, Group 5 cdDMRs (increasing glial, decreasing neuronal DNAm) were enriched in ASD genes from the SFARI Gene database (OR=5.7, FDR=4.1e-03), while Group 6 cdDMRs (decreasing glial, static neuronal DNAm) were enriched in ASD, syndromal neurodevelopmental disorder, and intellectual disability genes (all with OR>3.1 and FDR<1.9e-02). In contrast, a curated set of neurodegenerative disorder genes showed no enrichment for cdDMRs, perhaps reflecting lesser relevance of the first two decades of postnatal epigenomic remodeling to the etiology of those disorders.

In the non-CpG context, we found significant enrichment of both increasing and decreasing mCpH levels in genes associated with schizophrenia, ASD, and syndromal neurodevelopmental disorders (all with OR>2.1 and FDR<2.0e-02). CpH hypomethylation in neurons was also enriched in the neurodegenerative disease gene set (OR=2.6, FDR=3.7e-03). Finally, significantly increasing mCpH was depleted in genes associated with intellectual disability (OR=0.34, FDR=3.7e-06). While enrichment for conflicting mCpH patterns are at first curious given the overall negative association between mCpH and gene expression, outside of the context of DMRs, individual mCpH could be associated both positively and negatively with expression. Indeed, many genes, exons, and PSI events whose expression both positively and negatively associated with both mCpH and mCpG were also enriched in genes associated with schizophrenia, ASD, and syndromal neurodevelopmental disorders (all with OR>2.1 and FDR<2.5e-02; **Table S9**). Overall these results suggest that these examples of dynamic methylation and associated isoform switching may play a role in the development of higher cognitive functions during brain maturation associated with these diseases.

## Discussion

Here we have created a single-base resolution map of the dynamic DNAm landscape across the first two decades of postnatal human brain development in two cell type-enriched populations. Using FANS-derived samples, we were able to identify 40% more developmentally-regulated regions of changing DNAm than were identified in homogenate DNAm cortical data. We profiled specific features of the DNAm landscape including LMRs, UMRs, PMDs and DMVs and found that across features, neurons were typified by a general accumulation of mCpG. In the absence of complementary cell type-specific chromatin data, characterizing known DNAm features provided a more granular view of the potential functional genomic state in these regions than the available predictions derived from a few homogenate cortical samples. Particularly in studies using human postmortem brain, where tissue is often subjected to long postmortem intervals and low pH that degrades less stable epigenetic signatures, DNAm is a robust and durable marker that can be used to map the functional genomic terrain. These DNAm maps complement recently available epigenomic maps of different modalities generated on FANS-derived samples in the psychENCODE Consortium^27^.

We further parsed the general accumulation of neuronal DNAm into six trajectories of cell type-specific developmental patterns and found that neuronal mCpG progressively diverged from a shared landscape with glia and bulk prenatal cortex as the brain matured. Importantly, these diverging patterns were most striking during infancy through the first five years of postnatal life. The human brain experiences an explosion of synaptic connections during this time period, to nearly double the number found in the mature adult brain^28^. Although previous work has underscored this timeframe in terms of rapid DNAm accumulation^1^, this is the first work to refine DNAm patterns to reflect cell type-specific gain and loss of mCpG and mCpH within this critical window. By parsing these neuronal and glial DNAm patterns, we have highlighted epigenetically dynamic regions that may be contributing to the developmental processes such as synaptogenesis occuring during this timeframe that establish the foundation for fine-tuning connections throughout the remainder of brain maturation. These results provide a finely resolved depiction of epigenetic plasticity being greatest during this period of life, and support other evidence that environmental experience during these years may have an especially enduring impact on brain function^29^.

mCpH is unusually abundant in neurons compared to other cell types and appears to undergo trinucleotide-specific reprogramming during differentiation from ECSs^17^. While most mCpH in ESCs occurs in the CAG context, neuronal mCpH predominantly accumulates in the CAC context^17^. Here we elaborate on this relationship, showing that while both mCAG and mCAC aggregate in neurons as they mature and mCAG is gained faster than mCAC overall, mCAC accumulates proportionally faster in both neurons and glia over time. Interestingly, although neurons and glia contained mCpH in both trinucleotide contexts, mCAG was more likely to have higher levels in glia than neurons or be decreasing over development; indeed, genes containing decreasing mCAG but not mCAC were strongly associated with neuronal biological processes. mCpH trinucleotide context, therefore, may have as yet not well understood ramifications in brain development.

In terms of the relationship between mCpH and mCpG, we found that while neighboring mC (i.e., mCpG+mCpH) was not correlated genome-wide, mC was highly correlated within the cdDMRs despite local mCpH not being correlated. In other words, there was a convergence of levels of all contexts of methylation within the cdDMRs that was not detected genome-wide. mCpH also recapitulated the pattern seen in mCpG of diverging from a shared DNAm landscape with glia. Given that mCpH and mCpG have previously been shown to work in concert to recruit MECP2 binding to fine-tune gene expression^30^, it is sensible that levels of both contexts would perhaps reflect a shared functional role within putatively regulatory cdDMR sequence, since cdDMRs were also enriched for gene bodies and brain enhancer sequence. This work quantifies this correlation for the first time, a DNAm relationship unique to only a selection of cell types that includes neurons.

The identification of widespread association of mCpG and mCpH with expression and specific splicing events, particularly in neuronal genes enriched for neuropsychiatric diseases, highlights a potential novel role of mCpH and further expands the role of mCpG in the regulation of gene expression in neurons. Splicing is predominantly a co-transcriptional process influenced by changes in chromatin modifications and RNA binding proteins; the effects of DNAm on splicing decisions is not yet well studied^31^. Here we found thousands of associations between mC levels and gene, exon and PSI expression in developing postnatal neurons, particularly featuring many exons that are exclusively associated with mCpH. Although it is not possible to establish a causal role for mC in these data, these analyses, which are summarized in the provided website, can empower other researchers to explore the connection between DNAm and alternative isoform use, a phenomenon particularly prevalent in the developing brain that is often associated with disease^32^.

We also explored the relationship between DNAm patterns and genetic associations with various phenotypes and found both expected and surprising associations. We confirmed enrichment for heritability of brain traits generally in neurons, and heritability for schizophrenia, a disorder with strong neurodevelopmental underpinnings, specifically in genomic regions losing DNAm preferentially in neurons over early postnatal development (ie, Group 3 cdDMRs). This result emphasizes the critical nature of neuronal development and maturation in early establishment of pathological connectivity and function for this adult-onset disorder, as most DNAm loss--generally associated with increased activity of a gene or regulatory element--occurred within the first five postnatal years. Indeed, Group 3 cdDMRs were present in genes such as *GRIN1*, *SYN1* and *CAMK2A*, and others involved in establishing synapse organization and function, a hallmark of early postnatal brain development, implicating abnormal genetic regulation of neuronal connectivity in schizophrenia development. Interestingly, heritability for PTSD was significantly enriched in regions preferentially gaining DNAm in the non-neurons over development (Group 4 cdDMRs), regions associated with neural precursor cell proliferation and cell-cell connective properties (Figure 1C). This association, particularly given the small amount of sequence covered by Group 4 cdDMRs, could lead to fruitful insights into susceptibility for PTSD and warrants further study.

We also found enrichment of genes associated with rare variants implicated in psychiatric and neurodevelopmental disease in a variety of cell types and developmental trajectories, highlighting the the genomic boundaries, developmental timing and cellular context of epigenomic remodelling of regulatory elements or expressed features associated with known risk genes. Two examples of this are *HDAC4* and *CACNA1B,* genes associated with ASD in the SFARI Gene database. We identified a 566 bp Group 3 (decreasing neuronal, static glial DNAm) cdDMR within an intron of *HDAC4*, a calcium-sensitive transcriptional repressor, and a 3.6 kb Group 5 (decreasing neuronal, increasing glial DNAm) cdDMR within *CACNA1B*, a gene encoding a voltage-gated calcium channel subunit (**Figure 4 BC)**. Even though both genes are implicated in ASD and both cdDMRs are hypomethylated in neurons, the timing of loss of mCpG suggests that *CACNA1B* activity occurs earlier in postnatal development than *HDAC4*. Given that ASD onset is typically in early childhood, these risk genes may therefore have differing implications in the etiology of ASD. Another example is a 646 bp Group 6 (decreasing glial, static neuronal DNAm) cdDMR that overlaps the last intron and exon of *AKT3*, a serine/threonine-protein kinase gene. Although the *AKT3* /1q44 locus has been associated with schizophrenia risk, the mechanisms are not yet known given that *AKT3* is involved in many biological functions^25^. Interestingly, this cdDMR selectively lost mCpG in glia beginning in infancy, suggesting that *AKT3* activity in human DLPFC may be localized to glia beginning in infancy or earlier (**Figure 4 D**). This work provides the first *ex vivo* look at the DNAm dynamics within human neurons and glia and thus allows for the first examination of those parameters within the relevant organ, the brain.

Despite these insights, our data invoke several caveats. While NeuN based FANS greatly improves identifying developmental DNAm changes over homogenate data, designating NeuN+ and NeuN- samples as “neurons” and “glia” is not completely accurate in that NeuN- samples will include signal from unlabeled neurons and mask non-neuronal diversity. However, recent work assessing brain regional DNAm differences between NeuN+ and NeuN- found that NeuN- contributed comparatively marginal variability in DNAm compared to NeuN+, suggesting that neuronal methylomes are much more dynamic than non-neurons^20^. Likewise, while a percentage of the bases in the DMRs identified in this work have also been previously shown to be differentially methylated by neuronal subtypes whose unique methylomes are masked using NeuN-based FANS^5,14,33^, the proportion of these subtypes should be stable over postnatal development^34^. Future epigenomics studies however can improve on the resolution of our study by isolating more specific neuronal subpopulations to refine the cellular specificity of these neuronal methylation changes largely occurring in the first few years of life.

Another caveat is that WGBS does not allow for the discrimination between mC and hydroxymethyl-cytosines (hmC), an intermediary in the demethylation pathway. Previous work has shown^5^ that only a fraction of CpGs have measurable levels of hmC, suggesting that our results are not confounded. In the cited study^5^, hmCpG signal from homogenate cortex represented 10% of the hypermethylation found in excitatory neuron mCpG, suggesting that most of the mCpG signal in our neuronal data likely is true mCpG. The level of hmC in the non-CpG context remains controversial, with some studies not identifying hmCpH^5^ and others detecting low amounts (1% in FANS-derived human glutamatergic and 0.47% in GABAergic neurons)^35^. Further, FANS-derived human oligodendrocytes showed little hmC in the same study^35^. Overlap of our cdDMRs with DMRs between neurons and oligodendrocytes in a study of mC in FANS-derived human PFC samples^32^ showed that cell type differences primarily reflected true mC rather than hmC contamination in WGBS, while hmC DMRs between neuronal subtypes primarily were reflected in hypomethylated neuronal cdDMR groups and not the hypermethylated neuronal groups that would potentially include hmC signal contamination (**Figure S7B**). Future work, however, should more closely examine the contribution of hmC to the global hypermethylation seen during neurodevelopment.

By mapping the changing DNAm landscape over human postnatal neuronal and glial development, we have identified unique trajectories of DNAm change particularly dynamic during the first five years of life that show convergence between mCpG and mCpH, as well as associations between single mCpG and mCpH and alternative splicing. These patterns may also help illuminate the mechanisms through which psychological, neurological and psychiatric traits are developed by placing known genetic contribution in an epigenomic context.

## Author Contributions

- A.J.P.: Conceptualization, Formal Analysis, Investigation, Visualization, Writing – Original Draft Preparation, Writing – Review & Editing.
- L.C.-T.: Conceptualization, Formal Analysis, Visualization, Software, Writing – Original Draft Preparation, Writing – Review & Editing.
- N.A.I.: Software, Data Curation.
- W.X.: Investigation.
- E.E.B.: Formal Analysis, Writing – Review & Editing.
- J.H.S.: Supervision.
- R.T.: Investigation.
- L.M.: Investigation.
- Y.J.: Investigation, Supervision.
- T.M.H.: Data Curation, Resources, Writing – Review & Editing.
- J.E.K.: Data Curation, Resources, Writing – Review & Editing.
- D.R.W.: Conceptualization, Funding Acquisition, Supervision, Writing – Review & Editing.
- A.E.J.: Conceptualization, Formal Analysis, Funding Acquisition, Supervision, Writing – Original Draft Preparation, Writing – Review & Editing.

## Data availability

Raw and processed sequence data that support the findings of this study have been deposited at www.Synapse.org with the accession code: syn5842535.

## Code availability

Code is available through GitHub at: https://github.com/LieberInstitute/brain-epigenomics.

## Methods

### Postmortem brain tissue acquisition and processing

As previously described in Jaffe et al. (2016)^2^, post-mortem human brain tissue was obtained by autopsy primarily from the Offices of the Chief Medical Examiner of the District of Columbia, and of the Commonwealth of Virginia, Northern District, all with informed consent from the legal next of kin (protocol 90-M-0142 approved by the NIMH/NIH Institutional Review Board). Additional post-mortem prenatal, infant, child and adolescent brain tissue samples were provided by the National Institute of Child Health and Human Development Brain and Tissue Bank for Developmental Disorders (http://www.BTBank.org) under contracts NO1-HD-4-3368 and NO1-HD-4-3383. Postmortem human brain tissue was also provided by donation with informed consent of next of kin from the Office of the Chief Medical Examiner for the State of Maryland (under Protocol No. 12-24 from the State of Maryland Department of Health and Mental Hygiene) and from the Office of the Medical Examiner, Department of Pathology, Homer Stryker, M.D. School of Medicine (under Protocol No. from the Western Institute Review Board). The Institutional Review Board of the University of Maryland at Baltimore and the State of Maryland approved the protocol, and the tissue was donated to the Lieber Institute for Brain Development under the terms of a Material Transfer Agreement. Clinical characterization, diagnoses, and macro- and microscopic neuropathological examinations were performed on all samples using a standardized paradigm, and subjects with evidence of macro- or microscopic neuropathology were excluded, as were all subjects with any psychiatric diagnoses. Details of tissue acquisition, handling, processing, dissection, clinical characterization, diagnoses, neuropathological examinations, and quality control measures have also been further described previously^36^. Homogenate postmortem tissue of the prefrontal cortex (dorsolateral prefrontal cortex, DLPFC, BA46/9) was obtained from all subjects.

### Fluorescence activated nuclei sorting

Nuclei were isolated from 100-300mg of pulverized DLPFC tissue using dounce homogenization followed by ultracentrifugation over a sucrose density gradient. Homogenization was performed on ice in 5mL lysis buffer [0.32M sucrose, 3mM magnesium acetate, 5mM calcium chloride, 5mM EDTA (ph 8.0), 10mM Tris-HCl (pH 8.0), 0.1% Triton X-100] and the resulting homogenate was layered over 38mL sucrose buffer [1.8M sucrose, 3mM magnesium acetate, 10mM Tris-HCl (pH 8.0)] and centrifuged at 139,800 x g for 2 hours at 4°C. Cellular debris and lysis and sucrose buffers were removed and the pelleted nuclei were resuspended in 500uL PBS. Nuclei were then labeled in a solution of anti-NeuN antibody conjugated to Alexa Fluor 488 (A60, Millipore, 1/1000) and 0.1% BSA, rocking for 30 minutes at 4°C, followed by the addition of DAPI. Nuclei sorting was performed at the Johns Hopkins School of Public Health Flow Cytometry Core with a MoFlo Legacy (Beckman Coulter) using Summit (version 4.3) software. The purity of sorted populations was determined to be >99% based on resorting NeuN+ and NeuN- populations through the same gates.

The identity of the NeuN+ and NeuN- populations as neuron-enriched and glia-enriched was confirmed by sequencing nuclear RNA from each population and determining that neuronal and glial biological processes were enriched in genes differentially expressed between the two groups (FDR<0.05; **Figure S1B**). Likewise, cell type marker gene expression patterns also corroborated the neuronal- and glial-enriched identities of NeuN+ and NeuN- samples (**Figure S1C**). The estimated proportion of neurons in homogenate DNA methylation data based on deconvolution using differentially methylated sites between NeuN+ and NeuN- samples was highly correlated with the empirical proportion of neurons (**Figure S1D**). Raw sorting data is shown in **Figure S16**.

### Nucleic acid extraction and RNA-seq library preparations

RNA was extracted from homogenate and sorted samples using TRIzol LS Reagent (Thermo Fisher Scientific) followed by the RNeasy MinElute Cleanup Kit (Qiagen). Genomic DNA extraction was performed using the DNeasy Blood and Tissue Kit (Qiagen). Bisulfite conversion of 600 ng genomic DNA was performed with the EZ DNA methylation kit (Zymo Research). RNA sequencing libraries were made with the TruSeq RNA Library Prep Kit (Illumina) and the RiboGone Low-Input Ribosomal RNA Removal Kit (Clontech). Library concentrations were measured using a Qubit 2.0 and library fragment sizes were measured using a Caliper Life Sciences LabChip GX. One hundred base-pair paired-end sequencing was run on an Illumina HiSeq 2000.

### Whole genome bisulfite library preparations and sequencing

Sequencing libraries were made with Illumina TruSeq DNA Methylation library preparation kits. Lambda (il) DNA sequence was spiked in at 1% concentration to assess bisulfite conversion efficiency. Library concentrations were measured using a NanoDrop and library fragment sizes were measured using an Agilent Bioanalyzer 2100. Libraries were spiked in with 10% PhiX to improve base calibration calls and subsequently sequenced on an Illumina X-Ten Platform with paired-end reads (2×150bp), targeting 30x coverage and Q30 > 70% read quality.

### Data processing/alignment

We sought to align the paired-end reads for each sample to the in-silico bisulfite-treated hg19 genome, which we created using the Bismark v0.15.0^37^ `bismark_genome_preparation` program. For each library of paired-end reads (one per sample), the following processing was performed (**Figure S17**):

- FastQC v0.11.4, to assess read quality, presence of adapter sequence, and overrepresented sequences.
- Trimmomatic v0.35^38^ to trim low quality and adapter-containing portions of the reads, with the following parameters: `PE -threads 12 -phred33 ILLUMINACLIP:/Trimmomatic-0.35/adapters/TruSeq3-PE.fa:2:30:10:1 LEADING:3 TRAILING:3 SLIDINGWINDOW:4:15 MINLEN:75`. This resulted in three sub-libraries of reads per sample: one paired-end sub-library, and then two single-end sub-libraries where the other corresponding paired read was trimmed to a length below the defined threshold.
- FastQC v0.11.4, on each of the three sub-libraries, to assess the improvement in read quality and adapter content following trimming.
- FLASh v1.2.11^39^, to merge the paired-end sub-library reads into longer single-end reads, as reads that overlapped around CpGs and CpHs might bias or at the least double-count the DNAm estimates. Furthermore Bismark^37^ could only be run on single- or paired-end reads and not a combination of both. This therefore further split the paired-end sub-library into three sub-libraries: the subset of paired-end reads that were merged into longer single-end reads, and then left and right single-end reads that could not be merged.
- Bismark v0.15.0^37^, to align each of the five now-single-end sub-libraries (left-trimmed, right-trimmed, FLASh-merged, FLASh-left-unmerged, and FLASh-right-unmerged) to the bisulfite-converted hg19 genome using bowtie2^40^ and the --non-directional argument.
- Resulting alignment (BAM) files across five sub-libraries were merged, sorted, and indexed using samtools v1.3^41^ to produce one large/merged BAM file per sample.
- Alignments with evidence of duplication were removed using the MarkDuplicates program in Picard tools v1.141, which systematically appeared to be localized to low complexity DNA sequence near centromeres.
- The Bismark^37^ `bismark_methylation_extractor` program was run on each post-duplicate-removed BAM file per sample to extract CpG and CpH DNAm levels.

We additionally aligned reads from each sample to the PhiX and Lambda (λ) genomes to compute quality control metrics related to sequencing and bisulfite conversion quality. The average percentage of reads mapping back to the λ genome was 1.32% and the average bisulfite conversion efficiency was 98.64%. The average bisulfite conversion efficiency was not associated (p>0.05) with cell type, age, cell type (adjusting for age), age (adjusting for cell type), and the interaction of them in the NeuN- (glia) and NeuN+ (neuron) samples as well as for age in the homogenate samples. The genome coverage decreased from an initial average of 43x to 10x across the processing stages as shown in **Figure S18 (coverage)**. For each of the processing stages, there was no significant difference between cell types (adjusting for age), age (adjusting for cell type) as well as the interaction between age and cell type (p Bonferroni>0.05). The genome coverage was extracted from the `FASTQC` reports and by using `bamcount` (https://github.com/BenLangmead/bamcount) v0.2.6.

### Identifying methylation features

We identified PMDs, UMRs and LMRs using the bioconductor package MethylSeekR (version 1.20.0). We obtained coverage information from the cleaned set of ~18 million CpGs by extracting coverage and methylation using the `getCoverage()` function from the bsseq bioconductor package^42^ v1.10.0. PMDs were called using `segmentPMDs()` and were visually inspected using `plotPMDSegmentation()`. To create a more stringent cutoff for PMDs, we filtered PMDs to those longer than 100 Kbp. We calculated the FDRs using `calculateFDRs()`, while masking the >100 Kbp PMDs, setting the m parameter to 0.5 and the FDR cutoff to 10. PMDs were further filtered to exclude overlaps with the UCSC “gap” database table from hg19 except for the gaps labeled heterochromatin. We calculated UMRs and LMRs using `segmentUMRsLMRs()`. DMVs were defined as UMRs in which pmeth was less than or equal to 0.15 and the width was greater than or equal to 5 Kbp.

### Identifying CpG differentially methylated regions (DMRs)

Using the bsseq^42^ Bioconductor package v1.10.0 we loaded the Bismark^37^ report files and filtered the CpG data to keep only the bases where all samples had a minimum coverage of 3 (18,664,892 number passed the filter). We smoothed the methylation values of the remaining CpGs using the `BSmooth()` function from bsseq with the `parallelBy=“sample”` option. To identify the age and cell type DMRs we used a model that adjusted for both covariates while for the interaction model we included an additional interaction covariate. We identified the DMRs using the bumphunter^43^ Bioconductor package v1.14.0 using the `maxGap=1000`, `B=250`, `nullMethod=“permutation”`, `smooth=FALSE` options, which tends to be conservative in DMR identification^43^. For the `cutoff` option we used 0.1 for the cell type DMRs adjusting for age, 0.005 for the age DMRs adjusting for cell type, and 0.009 for the age and cell type interaction DMRs. These first two parameter cutoffs correspond to 10% minimum DNAm differences between neurons and glia and 5% change in DNAm per decade of life across cell types, and were chosen based on functionally-relevant change in DNAm. The cutoff for the interaction model (cdDMRs) was based on selecting the equivalent percentile of change from the overall age model (86th percentile) - this percentile-based cutoff was in line with recommendations for selecting cutoffs for statistical models with less clear biological interpretations^43^. We used a family wise error rate (FWER) threshold of 5% to determine the DMRs: `fwer` output from the `bumphunter()` function. A small subset of DMRs involved a single CpG, which arises from having a more significant area (length times effect size) than any DMRs identified in null permuted data. All DMRs showed less than 10% median percent absolute bias to technical and biological covariates as shown in **Figure S19 (sensitivity)**.

### cdDMR processing

For the “interaction” DMRs with FWER<5% (cdDMRs) we extracted the methylation values from the glial and neuronal samples using bsseq^42^ v1.13.9 and then computed a mean methylation value per DMR for each cell type. We also calculated the mean interaction coefficient for each DMR across all the cytosines in the DMR by cell type. Using the mean coefficients by cell type we clustered the interaction DMRs using the `kmeans()` function with `centers=6`, `nstart=100` options. We chose `centers=6` based on biological interpretability of the results and because k=6 results in an optimal AIC for clusters computed with mean centered and scaled data. For each cytosine in the cdDMRs we calculated the t-statistic and coefficient for age explaining differences in methylation adjusted for cell type: `~ age + cell type`. For each DMR we computed the mean age coefficient by cell type and then calculated the median absolute coefficient across all cdDMRs. The neuronal / glia ratio is 1.5 for such median absolute age effects across the cdDMRs.

### Comparing homogenate vs. cell type-specific WGBS

We first filtered CpGs to those with coverage in all 55 postnatal samples. For homogenate samples, we used `lmFit()` and `ebayes()` from the limma^44^ Bioconductor package v3.34.5 to assess age-associated changes to DNAm levels with the linear model `~ Age`. For the cell type-specific samples, we used a linear model `~ Age * Cell Type` to assess cell type, overall age, and age in a cell type changes to DNAm levels. We subset CpGs to those that were significantly differentially methylated by age in homogenate samples (p<1×10^−4^) and plotted the coefficients for each CpG in **<u>Figure S2A-D</u>**. The relationship between each variable was quantified using Fisher’s exact test.

### Roadmap Epigenome enrichments

We computed the relative enrichments of different genomic regions using Epigenome Roadmap data^12^ by computing the proportion of bases in each of the 15 ChromHMM states for each of the cells and tissues provided by the Consortium. We compared the proportion of bases in each state within each candidate region set to the overall genome and computed the corresponding log2 enrichments between the regions and this genomic background. We compared DMR and mCpG-based methylation feature regions to all profiled cell types in the Consortium for these analyses.

### Assessing the contribution of neuronal subtypes

The percent of neuronal subtype-specific bases were calculated by reducing the total subtype-specific CpG-DMRs from Luo *et al.*^14^, reducing the bases in each group of cdDMRs and calculating the percent of cdDMR bases that intersected the merged subtype-specific CpG-DMR bases.

### Visualizing DMRs

Plots for the DMRs were made using bsseq^42^ v1.14.0, EnsDb.Hsapiens.v75 v2.99.0, and RColorBrewer v1.1-2. Genes within 20 kb of a DMR were retained for the plots. Genes and exons were included using the `annoTrack` argument and we used extend`=2000` for making the plots.

### CpH processing

Using Bismark v0.16.3^37^ we created report files using the methylation extractor program with the `CX_context` and `split_by_chromosome` options for the hg19 human genome in order to extract the methylation values for the CpHs. Then for each chromosome, using the bsseq^42^ Bioconductor package v1.10.0 we loaded the Bismark^37^ report files and added the c_context and trinucleotide_context information from Bismark using custom R code based on the bsseq internal code that uses the data.table package v1.10.4. After combining the results for each chromosome, we filtered the CpHs to keep only those where all samples had a minimum coverage of 5 (58,109,566 number passed the filter).

### Lister, et al. (2013) data processing

We downloaded the WGBS data from Lister *et al.* data^1^ with SRA accession SRP026048. We then processed and aligned the data following the same steps we used for our data. Using bsseq^42^ as in the CpH processing section, we extracted the methylation values from the Bismark^37^ report files and added the `c_context` and `trinucleotide_context` information per chromosome. We then merged the results for all the chromosomes retaining only the CpG and CpH positions we observed with a minimum coverage of 3 and 5 in our data, respectively. To assess the replication of our cell type DMR results we computed mean methylation differences across the CpG positions comparing neuron and non-neuron samples in the Lister et al. data^1^ using the `rowttests()` function from the genefilter package version 1.56.0. We then computed the mean difference for each of the DMRs and compared this mean difference against the DMR mean methylation difference derived from our data to derive the concordance and correlation between them. To assess the replication of our age DMR results we modeled age as a continuous variable and calculated the mean methylation difference per year for every CpG contained in the age DMRs using `lmFit()` function from limma. Finally, we compared the mean methylation difference in the Lister et al. data^1^ against the observed mean methylation difference for the age DMRs we derived.

### Homogenate data processing

We processed the prenatal and postnatal homogenate brain samples using the same procedure described in the *Data processing/alignment* section to produce Bismark^37^ report files. Then using bsseq^42^ we extracted the methylation values for the CpG positions observed in our postnatal samples in order to make them comparable to each other.

### Identification of differentially methylated positions

With the set of CpGs and CpHs with a minimum coverage 3 and 5 in all samples, respectively, and the same models for identifying the DMRs, we identified the differentially methylated positions (DMPs), keeping the CpGs and CpHs separate. For the CpHs we further filtered to keep only those where at least 5 samples had a methylation value greater than 0 (40,818,742, 70.2%). We used the limma Bioconductor package v3.30.13 for determining the DMPs by running functions `lmFit()` and `eBayes()` with default parameters and FDR<5%.

### Enrichments for HARs and enhancers

We calculated the enrichment of genomic segments overlapping cdDMRs and human accelerated regions (HARs) and enhancers^15^ using Fisher’s exact test. We calculated the overlap of methylation features (DMRs, mCpH, expression-associated cytosines) and HARs or enhancers with the entire set of CpG clusters used to identify the DMRs as background. We corrected for multiple testing using the false discovery rate (FDR).

### Stratified linkage disequilibrium score regression

GWAS summary statistics for 30 phenotypes (described in Anttila et al. and Rizzardi et al.)^20,21^ were downloaded from the sources listed in Table S6. We used LDSC (LD SCore) v1.0.0 to estimate the proportion of heritability captured in 16 sets of genomic DNAm features for each GWAS phenotype, including the DMRs and LMRs defined above in this work, non-differentially methylated CpG clusters (called above before the DMR analysis, excluding DMR sequence), central nervous system annotations included in the LDSC package (referred as CNS (LDSC)), and regions annotated as putatively regulatory in human brain using chromHMM (i.e., the union of regions annotated as “Bivalent Enhancer”, “Bivalent/Poised TSS”, “Genic enhancers”, “Flanking Active TSS”, “Active TSS”, “Strong transcription”, and “Enhancers” in the following tracks accessed using the AnnotationHub Bioconductor package (v2.14.5)^45^: AH46920, AH46921, AH46922, AH46923, AH46924, AH46925, AH46926, AH46927, AH46934, and AH46935).

We first converted the GWAS summary statistics into the `.sumstats` format using `munge_sumstats.py` and keeping only HapMap 3 SNPs (downloaded from https://data.broadinstitute.org/alkesgroup/LDSCORE/w_hm3.snplist.bz2) as described in the Partitioned Heritability LDSC tutorial. We made `.annot` files for each custom feature set based on the list of SNPs in the CNS cell type annotations provided in the LDSC package and estimated partitioned LD scores for each feature using 1000 Genomes plink files (downloaded from https://data.broadinstitute.org/alkesgroup/LDSCORE/1000G_Phase3_plinkfiles.tgz) using `ldsc.py`. We finally estimated the partitioned heritability for each feature-phenotype combination by adding each feature individually to the “baseline model” including 53 baseline annotations described in Finucane et al.^22^

### Enrichments for genes associated with brain disorders

We calculated the enrichment of genes overlapping different methylation features in gene sets described by Birnbaum et al.^24^ We measured what fraction of the genes in each set overlapped methylation features (ie, DMRs, mCpH, and expression-associated cytosines) with Fisher’s exact test using all expressed genes with Entrez IDs as background. We corrected for testing multiple disorder gene sets using the false discovery rate (FDR).

### RNA-seq data processing

Raw sequencing reads were mapped to the hg19/GRCh37 human reference genome with splice-aware aligner HISAT2 v2.0.4^46^. Feature-level quantification based on GENCODE release 25 (GRCh38.p7) annotation on hg19 coordinates was run on aligned reads using featureCounts (subread v1.5.0-p3)^47^. Using custom R code we processed the different feature counts and created RangedSummarizedExperiment objects using the SummarizedExperiment Bioconductor package v1.4.0.

### Percent spliced in (PSI) calculation

We calculated the percent spliced in using the SGSeq^48^ Bioconductor package v1.12.0 and the Gencode v25 annotation for the GRCh37 human reference genome (ftp.sanger.ac.uk/pub/gencode/Gencode_human/release_25/GRCh37_mapping/gencode.v25lift <u>37.annotation.gtf.gz</u>) from the BAM files generated by HISAT2. We used default arguments except for the function `analyzeVariants()` where we used a `min_denominator=10`.

### Differential expression between cell types

We combined the gene counts for the polyA+ and RiboZero sequencing protocols for the cell-sorted RNA-seq data: 3 NeuN+ and 3 NeuN- samples for a total of 12 RNA-seq sequencing runs. We calculated the library size normalization factors using `calcNormFactors()` from edgeR^49^ (v3.22.3) and identified the differentially expressed genes using `voom()`, `lmFit()` and `eBayes()` from limma^44,50^. We repeated this procedure for the exon counts.

### Gene ontology analyses

Gene ontology enrichment analyses were performed using clusterProfiler^51^ v3.6.0 using the options `pAdjustMethod=“BH”`, `pvalueCutoff=0.1`, and `qvalueCutoff=0.05` on the Entrez IDs for each expression feature for the BP, CC and MF GO ontologies. Only cytosines or DMRs overlapping genes were included.

### Methylation vs expression associations

With the sorted RNA-seq data, we computed the average gene expression per age group (infant, child, teen) and cell type (six groups), and correlated these values to average DNAm levels in the same groups in both the CpG and CpH contexts at the gene promoter and body, exon (500bp window), and splice junction (50bp into each intron) levels. With the RangedSummarizedExperiment objects with the RNA-seq polyA homogenate data and the bsseq objects with the CpG and CpH data we determined which CpG and CpH positions explained changes in expression (RPKM) at the gene or exon level as well as in percent spliced in (PSI). We retained only the expression and PSI data from the postnatal samples and matched them by brain identifier to the neuronal methylation data with a final sample size of 22. We filtered lowly expressed features using the `expression_cutoff()` function from the jaffelab package v0.99.18: mean RPKM>0.22 mean for genes, 0.26 for exons. For genes and exons we transformed the expression values to log2(RPKM + 1) and extracted the raw PSI values. Using MatrixEQTL^52^ v2.2 (GitHub b9a9f01 patch) we then identified the methylation quantitative trait loci (QTL) for the CpG and CpH methylation data separately using the function `Matrix_eQTL_main()`function with options `pvOutputThreshold=0`, `pvOutputThreshold.cis=5e-4`, `useModel=modelLINEAR`, `cisDist=1000`. We identified marginal CpG associations near CpH associations by running MatrixEQTL again for the CpG in a +- 1kb window around the CpH positions with an association with expression at FDR <5% for each expression feature type using the same parameters as above except for `pvOutputThreshold.cis=0.01`. We filtered the associations to retain only those having at least 11 samples with non-zero methylation and 11 samples with non-one methylation values to remove extreme cases. We further restricted the results to protein coding genes and dropped any with infinite t-statistics. To assess whether age confounds the relationship between methylation and expression, we used a multiple linear regression model adjusting for age and checked if the methylation coefficient was still FDR<5%. Venn diagrams in **Figure 3** were made with the VennDiagram package v1.6.18. For the gene and PSI associations we used the gene ID to check if it was present in the 3,473 differentially expressed genes from the sorted RNA-seq data (described above) with higher expression in neurons and the top 5,000 DE genes with higher expression in glia at FDR<5%; similarly we did so for exons and the top 5,000 DE exons (FDR<5%) with higher expression in each cell type. The LIBD WBGS Expression explorer at https://jhubiostatistics.shinyapps.io/wgbsExprs/ was made using the bsseq^42^ v1.14.0, DT v0.4, SGSeq^48^ v1.12.0 and shiny v1.0.5 R packages.

### Global autocorrelation

Using CpG and CpH positions with a minimum coverage of 3 and 5 respectively for all samples we calculated the autocorrelation for the methylation levels for the CpGs, the CpHs, the CpHs with a CHG trinucleotide context, or the CpHs with a CHH trinucleotide context. For each of the sets, we grouped the positions using derfinder v1.12.0 into groups by a maximum distance of 1 kb. Only those groups with at least 5 Cs were further considered. For each sample we then calculated the autocorrelation using the `acf()` function with `lag.max=4` in parallel for each chromosome using BiocParallel v1.12.0. For each cluster of cytosines we calculated the mean across the neuronal (NeuN+) and the glia (NeuN-) samples at each autocorrelation lag. After combining and tidying the results, we visualized the global auto correlation using ggplot2 v2.2.1. We repeated this same analysis for the Lister et al. data^1^.

### Autocorrelation within DMRs

Similar to the global autocorrelation, we extracted the methylation values at CpGs with a minimum coverage of 3 and the CpHs with a minimum coverage of 5 that were within each of the sets of DMRs (age, cell type or interaction). We then computed the autocorrelation for DMRs with a least 5 different cytosines using the acf() function with a lag.max=4 and calculated the mean auto-correlation among the neuronal and glial samples. The lag is proportional to the genomic distance as shown in **Figure S20 (lag and distance)**.

---

**Figure S1:**
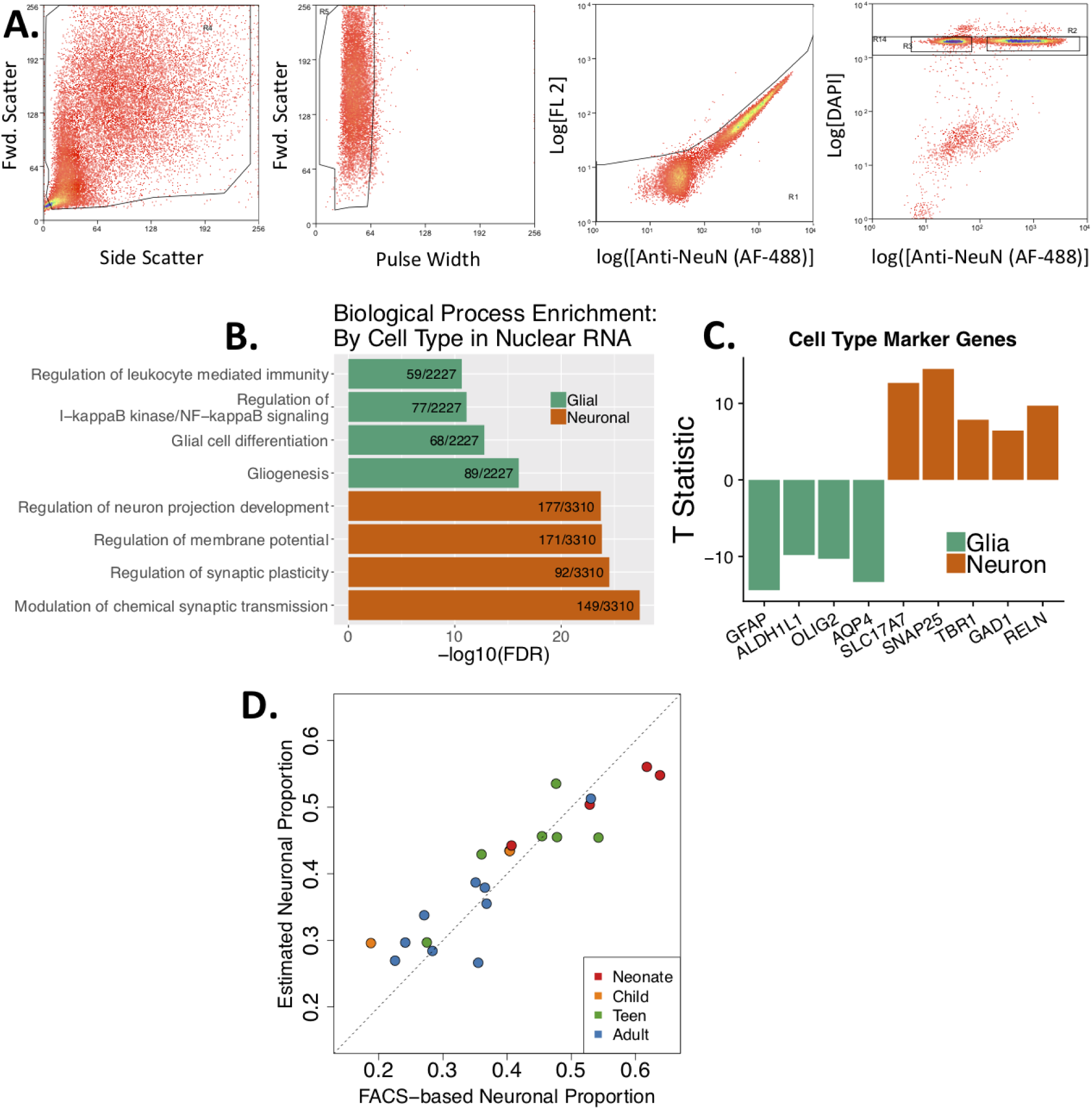
Confirmation of neuronal- and glial-enriched identity of NeuN+ and NeuN- samples. **(A)** Representative gating strategy from fluorescence-activated nuclear sorting by NeuN antibody. Debris is first reduced by selecting events based on forward scatter and side scatter, then aggregates are reduced by measuring pulse width. Autofluorescent events are discarded by measuring true Alexa fluor-488 signal compared to signal in the FL2 channel. Finally, singlets are determined by DAPI staining, and Alexa fluor-488 positive and negative events are collected. **(B)** Top biological processes enriched in genes significantly differentially expressed in nuclear RNA from NeuN+ (labeled “Neurons”) and NeuN- (labeled “Glia”) at FDR<0.05. **(C)** T statistics for marker genes for neuronal and non-neuronal identity show that the genes are differentially expressed in NeuN+ and NeuN- nuclear RNA (FDR<0.05). **(D)** The estimated proportion of neurons in homogenate WGBS samples based on statistical deconvolution using differentially methylated cytosines between NeuN+ and NeuN- samples is highly correlated with the empirically-derived proportion of NeuN+ events.

**Figure S2:**
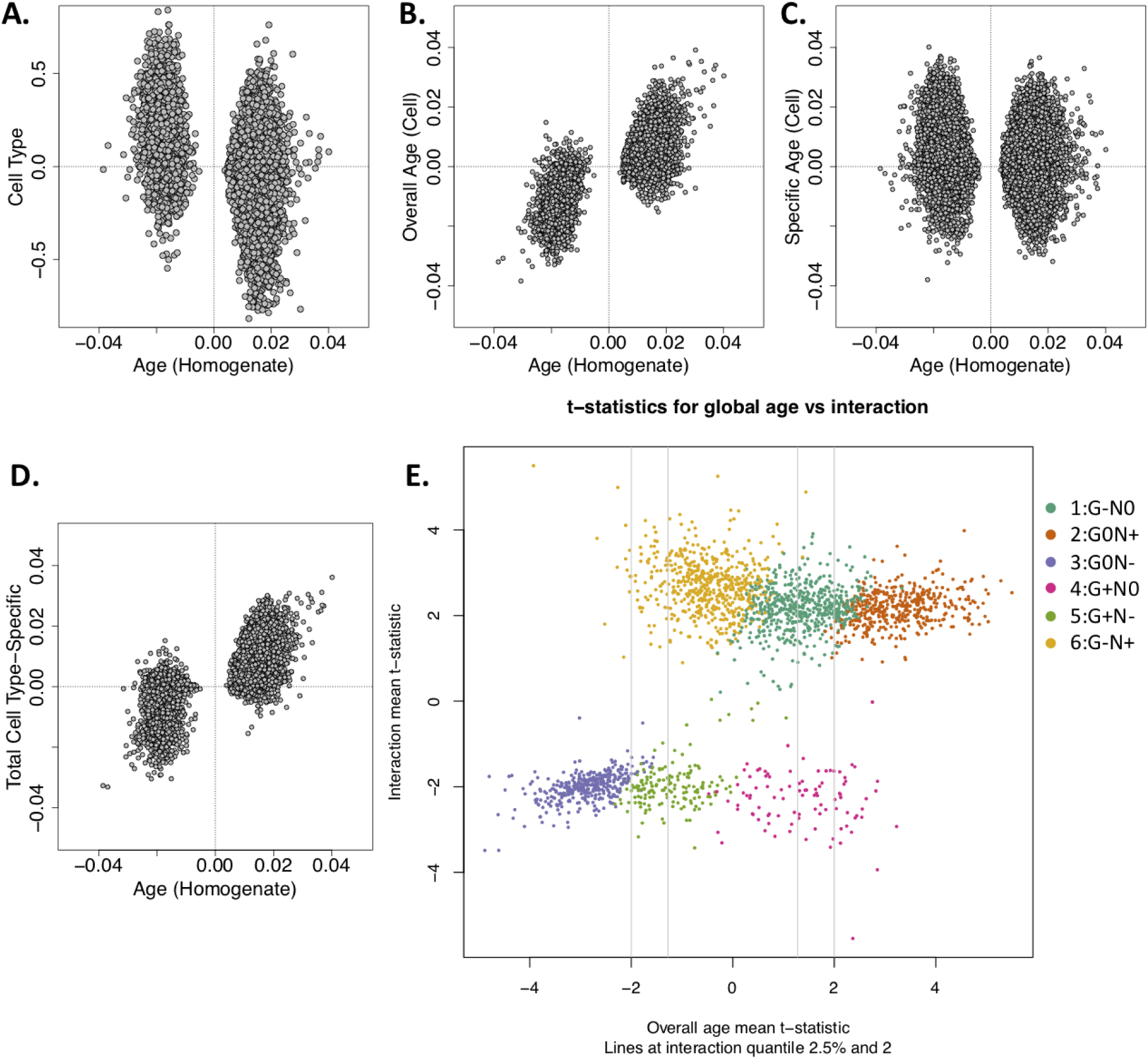
Detecting developmental changes in homogenate vs. cell type-specific DNAm data. **(A)** Developmental age effect coefficients of individual CpGs as measured in homogenate samples compared against cell type effect coefficients in cell type-specific samples after adjusting for age. **(B)** Developmental age effect coefficients of individual CpGs as measured in homogenate samples compared against the developmental effect adjusting for cell type in cell type-specific samples. **(C)** Developmental age effect coefficients as measured in homogenate samples compared against the cumulative age and cell type interaction effect coefficients in cell type-specific samples at the CpG level. **(D)** Developmental age effect coefficients as measured in homogenate samples compared against the estimated cell type-specific age effects for cell type-specific samples at the CpG level. **(E)** Overall age mean t-statistic for the cdDMRs (x-axis) against the mean interaction t-statistic (y-axis) with the 2.5% quantile from the y-axis shown on the x-axis. Colors are the same as those from Figure 1B: teal=Group 1 (decreasing glial methylation, increasing neuronal methylation), orange=Group 2 (static glial methylation, increasing neuronal methylation), purple=Group 3 (static glial methylation, decreasing neuronal methylation), pink=Group 4 (increasing glial methylation, static neuronal methylation), green=Group 5 (increasing glial methylation, decreasing neuronal methylation), gold=Group 6 (decreasing glial methylation, static neuronal methylation).

**Figure S3:**
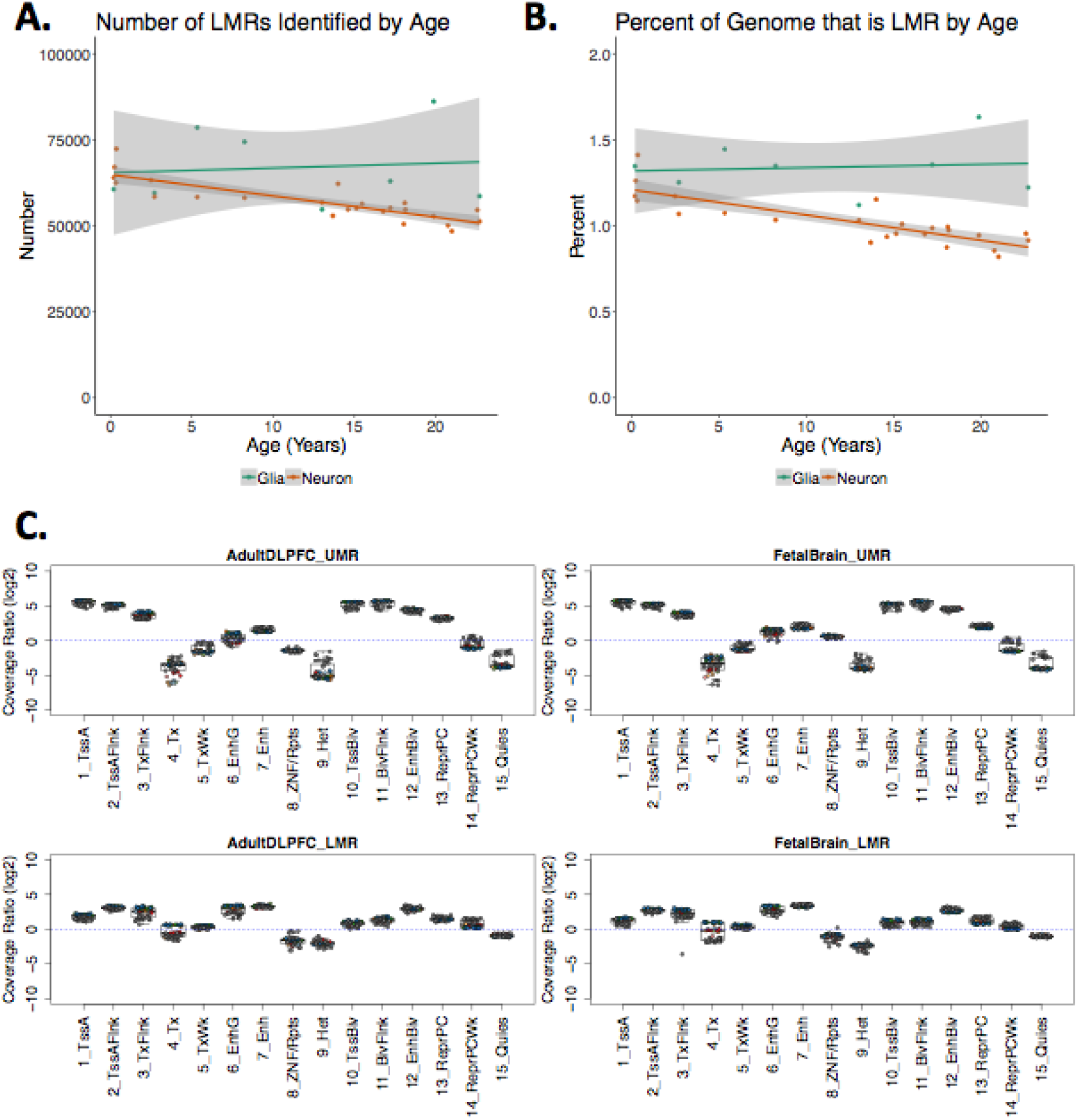
Unmethylated Regions (UMRs) and Low-methylated regions (LMRs). **(A)** Number of LMRs and **(B) p**ercent of the genome covered by LMRs by age stratified by cell type. Shading indicates the standard error of the linear model. **(C)** Roadmap Epigenomics Consortium predicted chromatin-enriched states for LMRs and UMRs as defined in adult DPLFC and fetal brain. log2(Coverage Ratio) represents the enrichment of the proportion of bases within LMRs or UMRs in a chromatin state compared to the rest of the genome.

**Figure S4:**
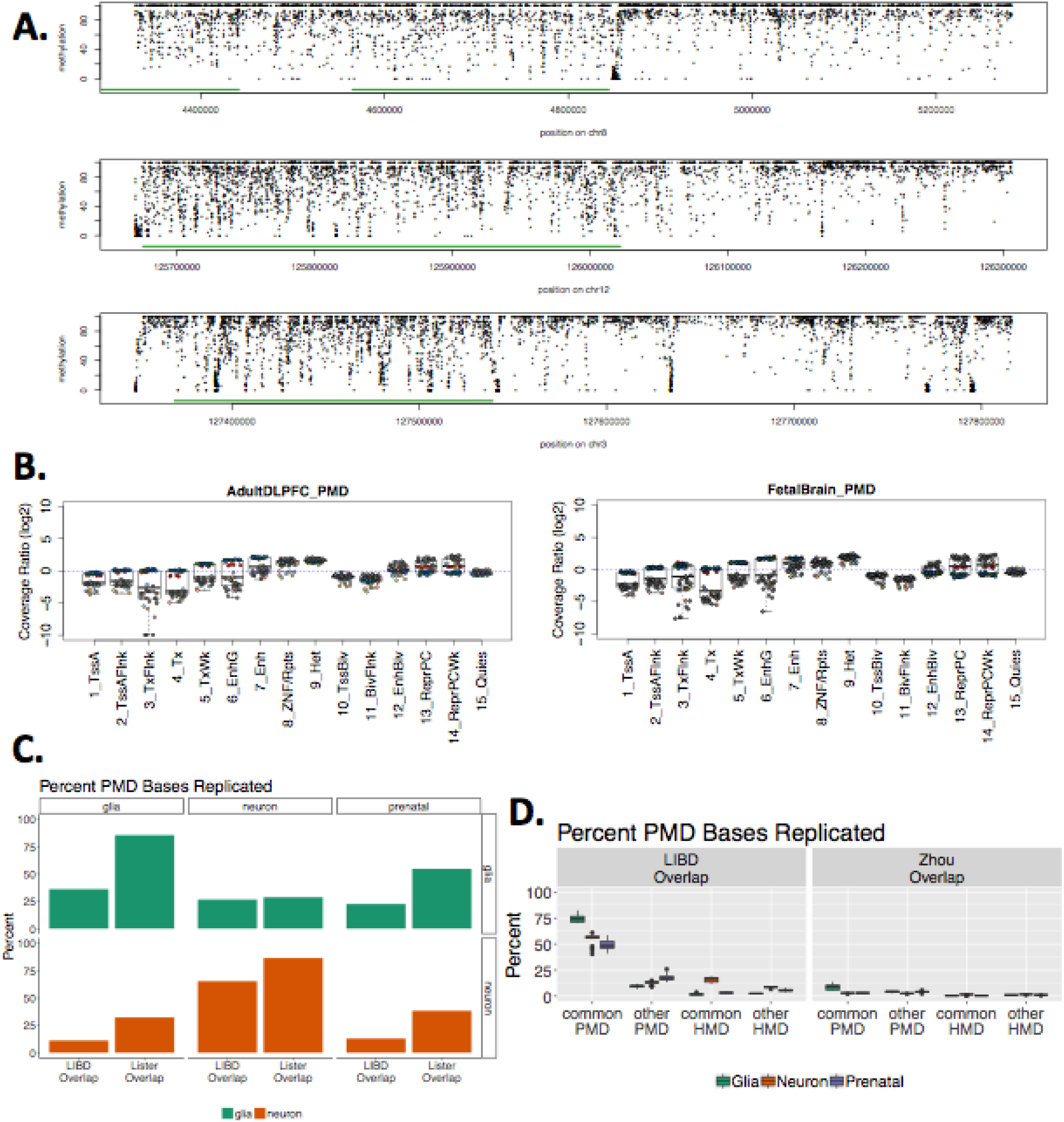
Partially methylated domains (PMDs). **(A)** Example PMDs on chromosomes 8, 12, and 3. PMDs are highlighted in green. **(B)** Roadmap Epigenomics Consortium chromatin state enrichment for PMDs. log2(Coverage Ratio) represents the enrichment of the proportion of bases within PMDs in a state compared to the rest of the genome. **(C)** Percent of PMD base-pairs that are replicated in the Lister *et al.* (2013) FANS samples^1^. Rows represent Lister *et al.* (2013) glia and neurons, and columns represent the glia, neurons and prenatal samples from this paper (LIBD). Each bar represents the percent of PMD bases shared between that quadrant’s cell types, using either total LIBD or total Lister PMD bases as the denominator. **(D)** Percent of PMD base pairs per sample that are replicated in either common or unique PMDs and high methylated domains (HMDs) identified in Zhou *et al.^13^* Samples are colored based on whether the sample was postnatal neuron, postnatal glia, or bulk prenatal cortex.

**Figure S5:**
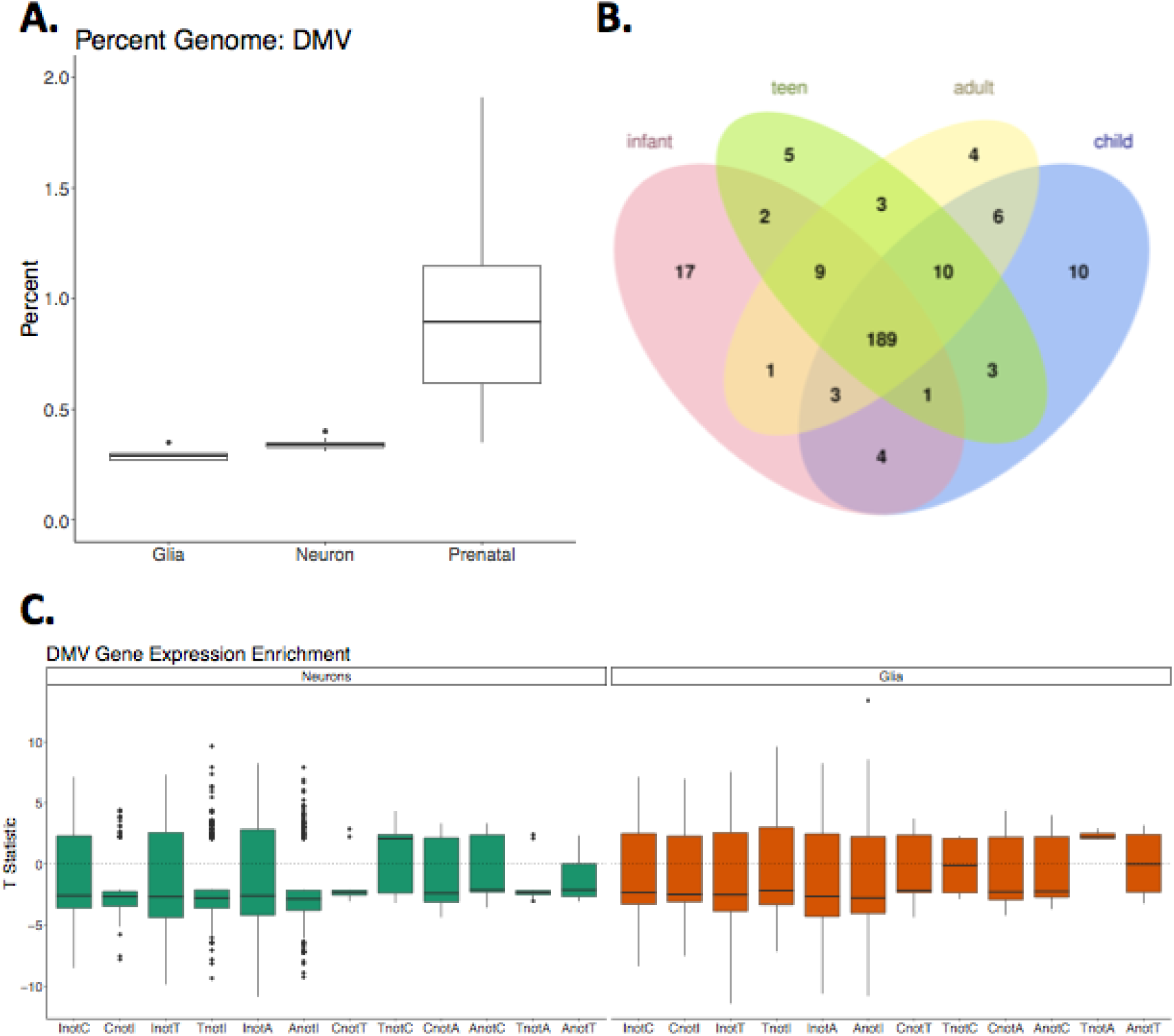
DNA methylation valleys (DMVs). **(A)** Percent of the genome covered by DMVs in postnatal glia and neurons and bulk prenatal cortex. **(B)** Overlap of transcription factor (TF) genes within DMVs by age in neurons. **(C)** Expression enrichment between age groups in TF genes excluded from DMVs in one group but not the other in neurons and glia. A negative T-statistic signifies greater expression in the age group in which the gene is not in a DMV. I=Infant (0-1 years); C=Child (1-10 years), T=Teen (11-17 years) and A=Adult (18+ years).

**Figure S6:**
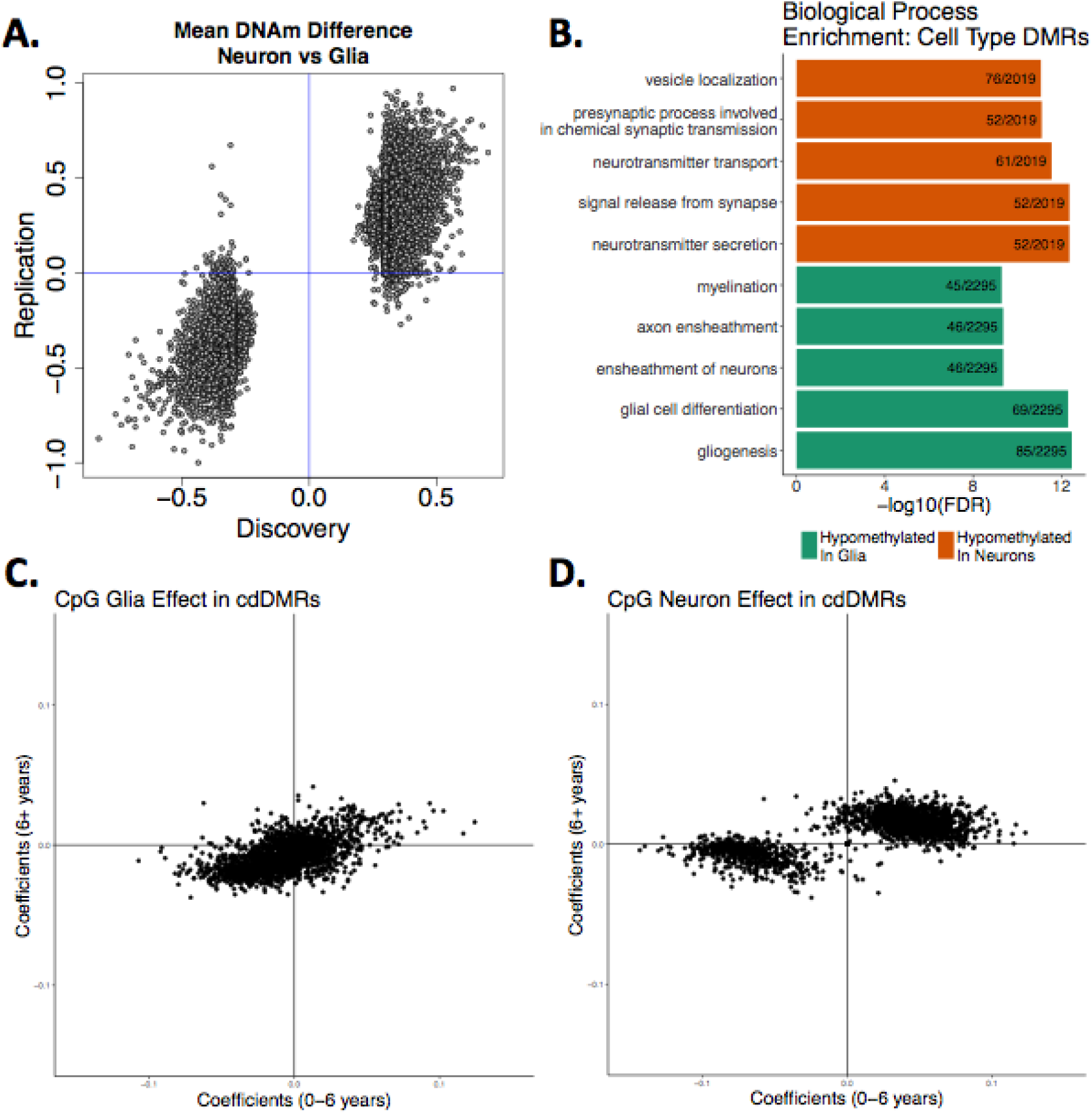
Differentially methylated regions (DMRs). **(A)** Replication of cell type DNAm differences at the CpG level between neurons and glia (NeuN+ and NeuN- samples) in our and Lister *et al.* data^1^. X and Y axes are linear model coefficients. (**B)** Enriched biological process ontology terms in the DMRs by cell type. The number of genes within each ontology group that overlaps a cell type DMR is listed. (**C)** Coefficients of linear regression on the mean mCpG level per cdDMR in samples younger than 6 years and older. **(D)** Coefficients of linear regression on the mean mCpG level per cdDMR in samples younger than 6 years and older.

**Figure S7:**
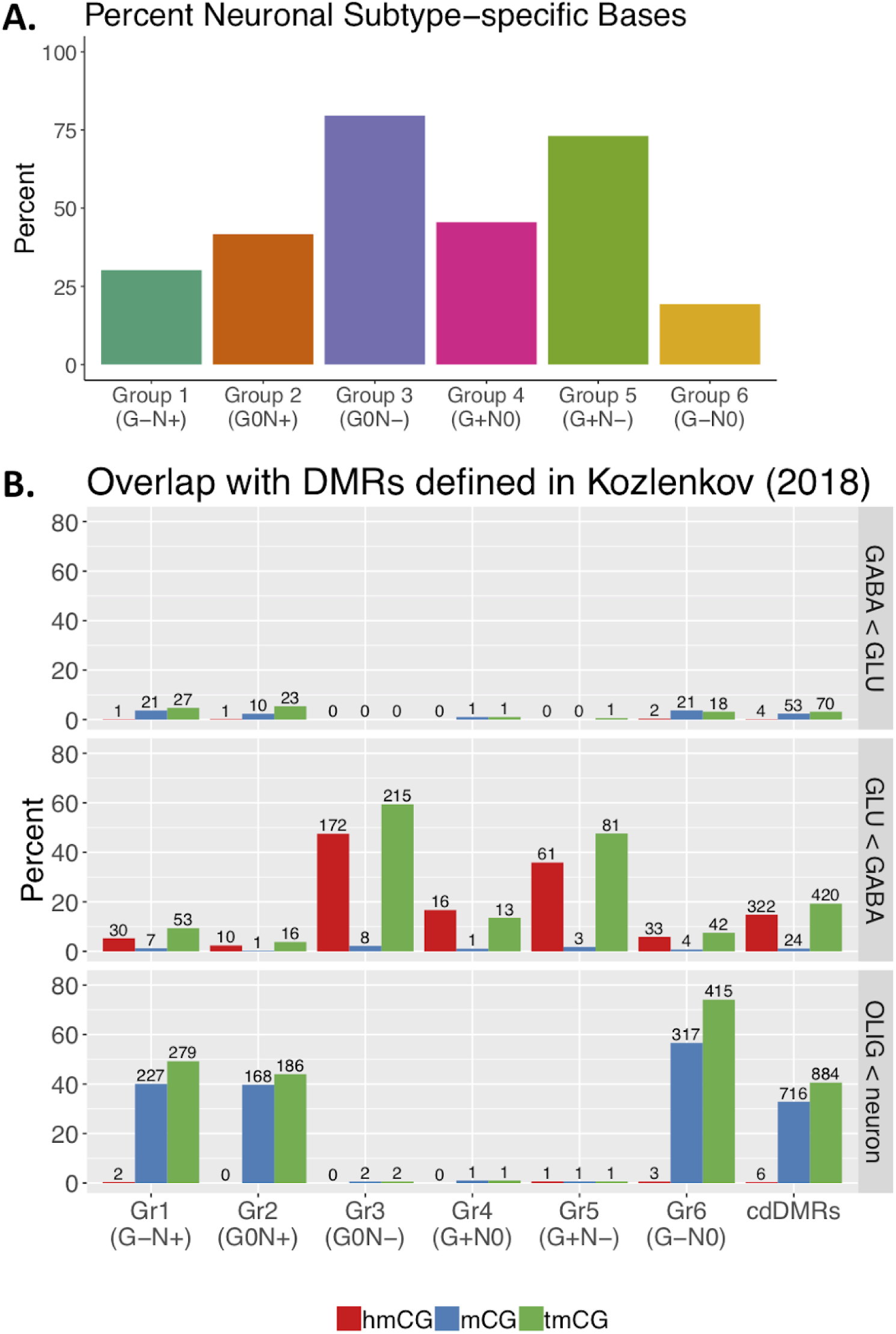
Cell type-specific developmental DMRs and neuronal subtype methylation and hydroxymethylation. **(A)** Percent of bases within each group of cdDMRs as defined in Figure 1B that are differentially methylated by neuronal subtypes in Luo *et al.^14^*. **(B)** Percent of cdDMRs within each of the 6 groups as defined in Figure 1B that overlap DMRs of methylcytosine (mC captured using oxidative bisulfite sequencing), hydroxymethylcytosine (hmC), or total cytosine methylation (tmC or mC+hmC, captured using standard bisulfite sequencing) as defined by Kozlenkov *et al.^50^*. The plot is stratified by whether the Kozlenkov *et al.* (2018) DMRs are more highly methylated in GABAergic or glutamatergic neurons, or in overall neurons compared to oligodendrocytes. The number of overlapping cdDMRs are listed above each bar.

**Figure S8:**
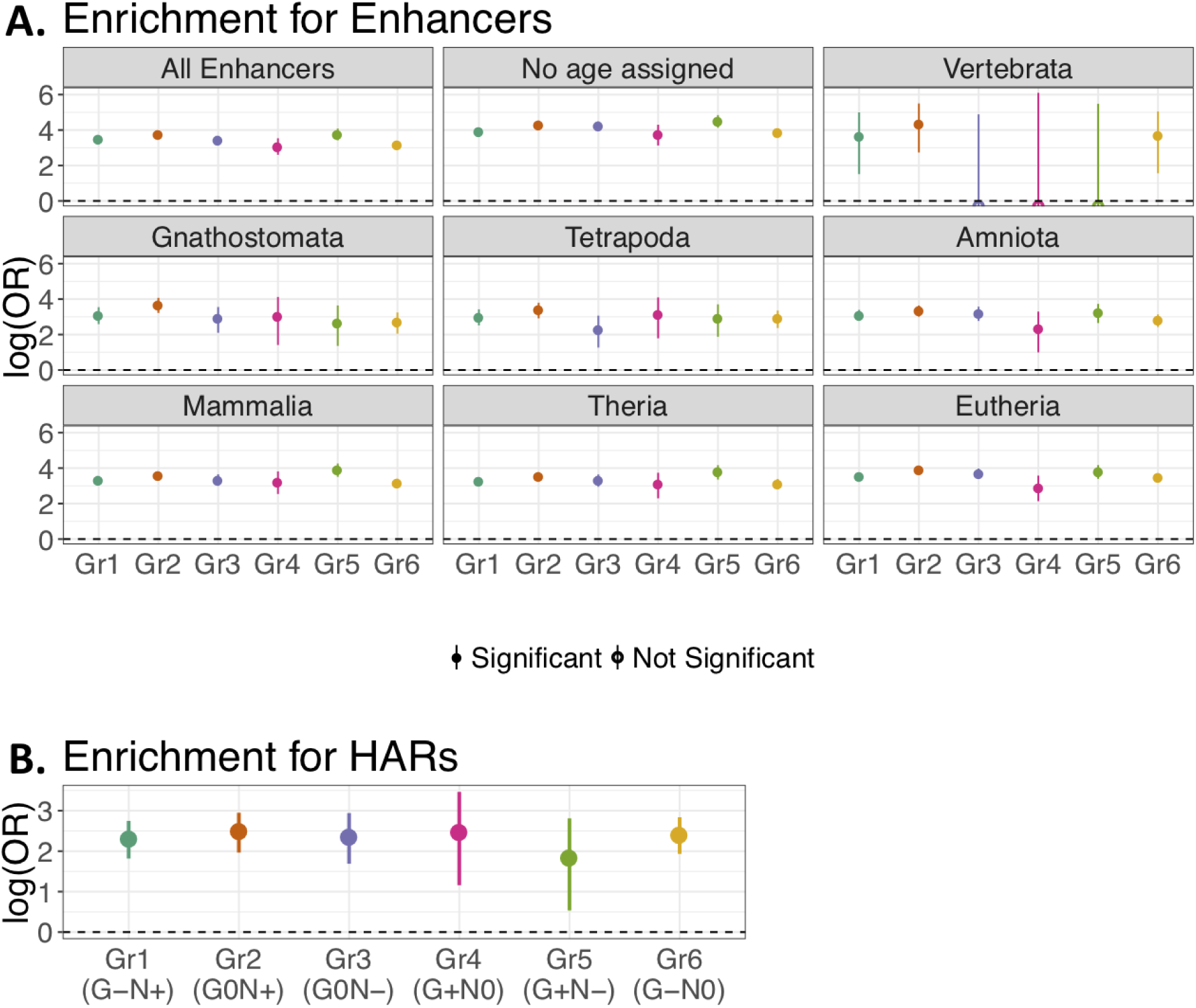
cdDMR overlap of human brain developmental enhancers and Human Accelerated Regions (HARs). **(A)** Enrichment for enhancers in the six clusters of cdDMRs from Figure 1B at the CpG-level. **(B)** Enrichment for human accelerated regions (HARs) in the six clusters of cdDMRs from Figure 1B at the CpG-level. Filled in circles indicate FDR<0.05, and error bars are the 95% confidence interval for the log(odds ratio).

**Figure S9:**
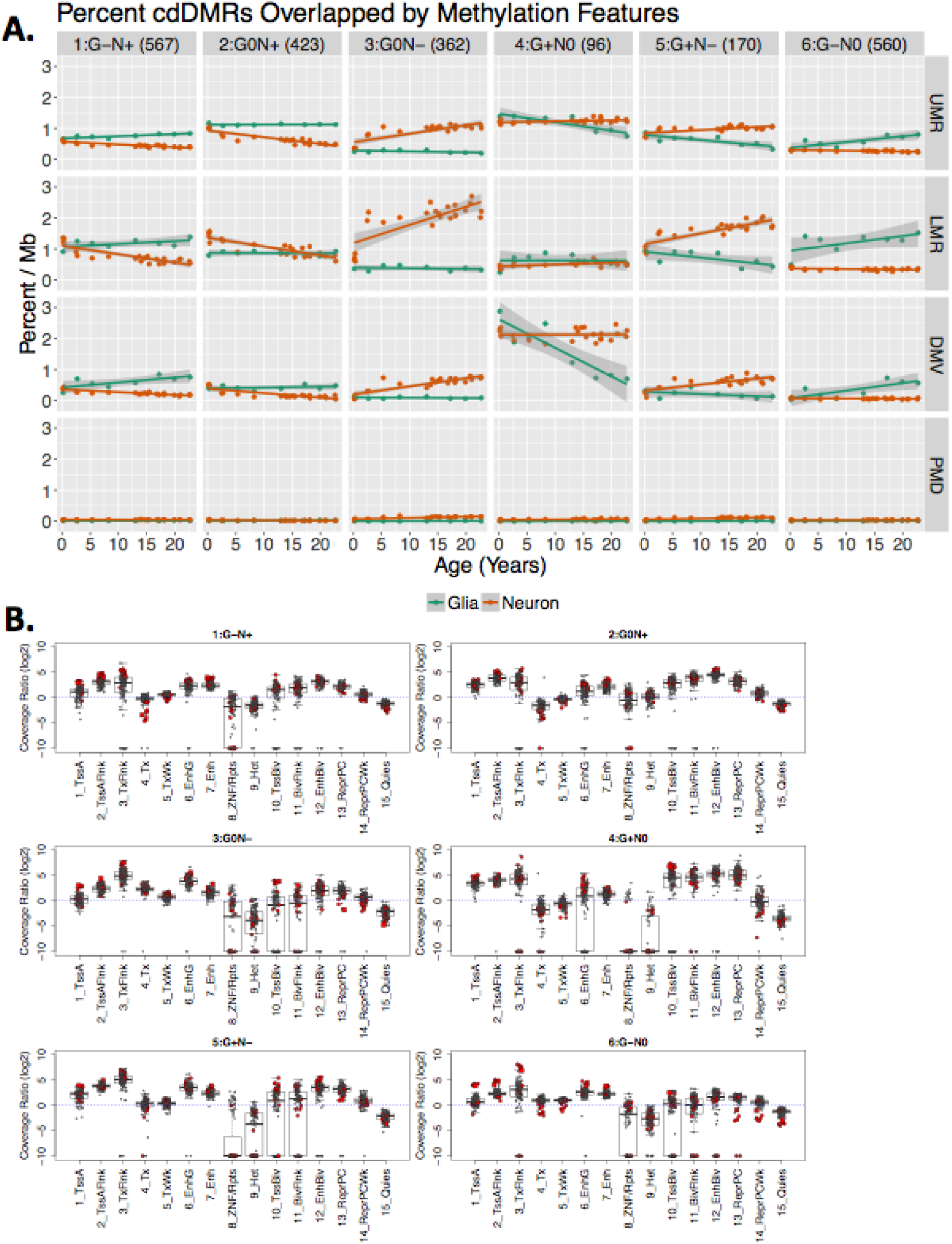
Cell type-specific, developmentally dynamic DMRs (cdDMRs) and epigenetic states. **(A)** Percent of cdDMRs by the k-mean groups from Figure 1B overlapped by UMR, LMR, DMV and PMD sequence across age. Lines show linear regression and shading indicates the standard error. **(B)** Roadmap Epigenomics Consortium enriched chromatin states for the six clusters of cdDMRs from Figure 1B at the CpG-level. log2(Coverage Ratio) represents the enrichment of the proportion of bases within each cdDMR group in a chromatin state compared to the rest of the genome.

**Figure S10:**
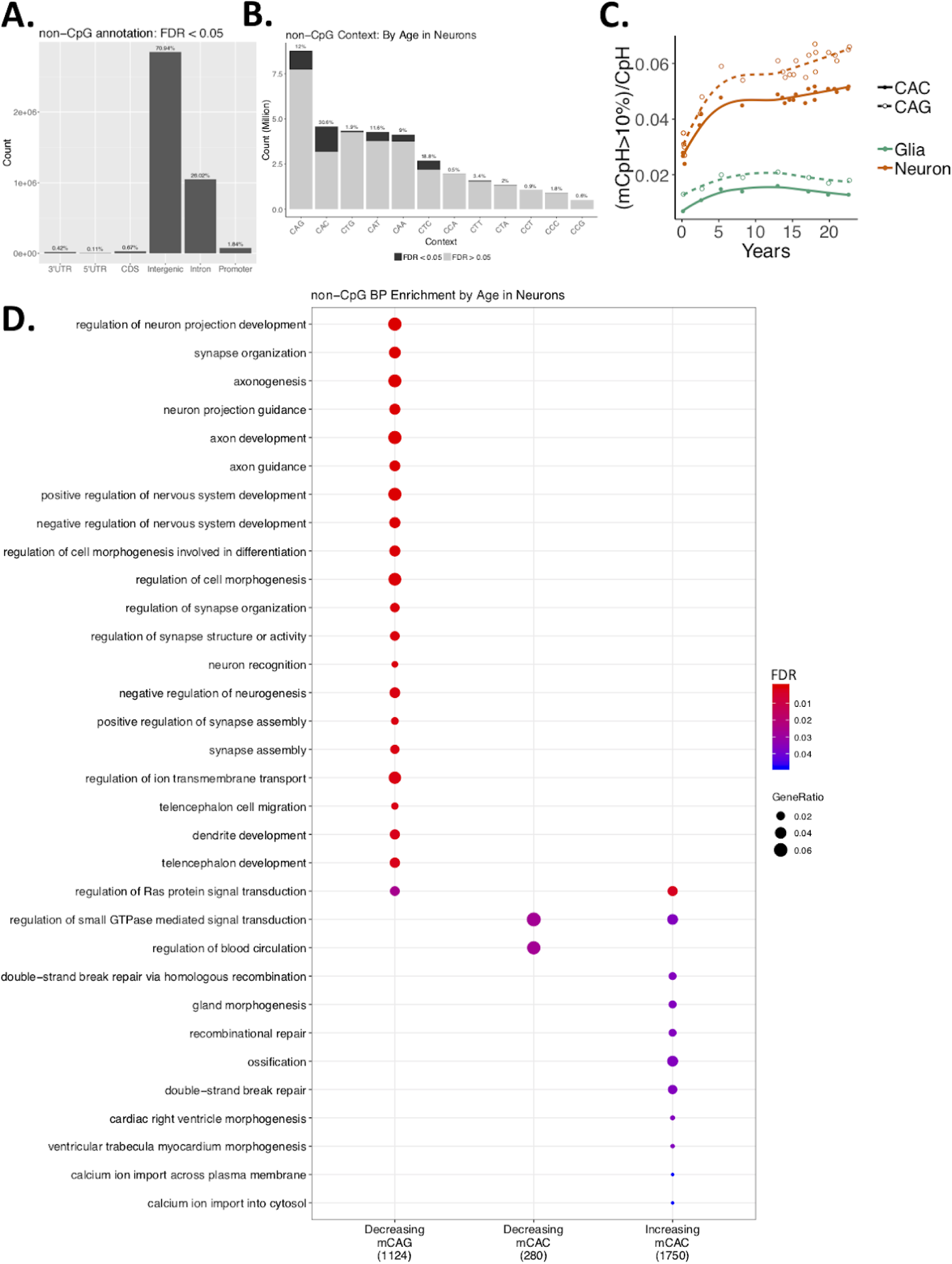
CpH methylation distribution, levels and context-specific biological process ontology. **(A)** Number of differentially methylated CpHs by cell type (FDR<5%) falling in different genomic annotations across the genome. Annotation was prioritized CDS > 5’UTR > 3’UTR > Intron > Promoter > Intergenic. **(B)** Breakdown of measured CpH by trinucleotide context. Differentially methylated mCpH by age in neurons (FDR<0.05) are colored in dark gray. **(C)** Accumulation of mCpH by trinucleotide context over development, stratified by cell type. **(D)** The top 20 biological process ontology terms enriched for genes exclusively overlapping CAG and CAC sites with significantly increasing or decreasing methylation levels in neurons over postnatal development (FDR<0.05).

**Figure S11:**
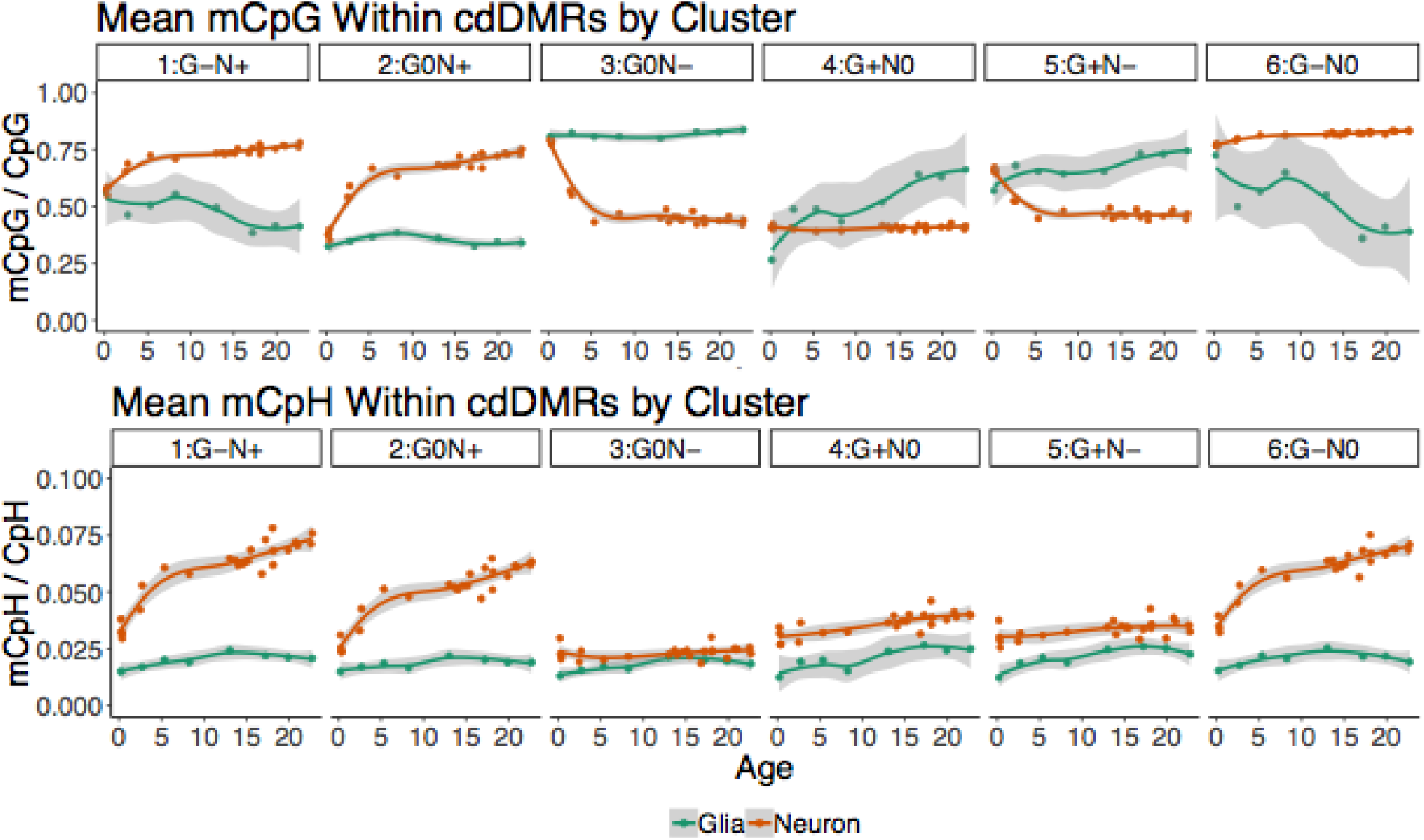
Trajectories of methylation accumulation in cdDMR groups. Mean mCpH and mCpG within the cdDMRs by cluster from Figure 1B across development. Loess line and standard error shading are depicted.

**Figure S12:**
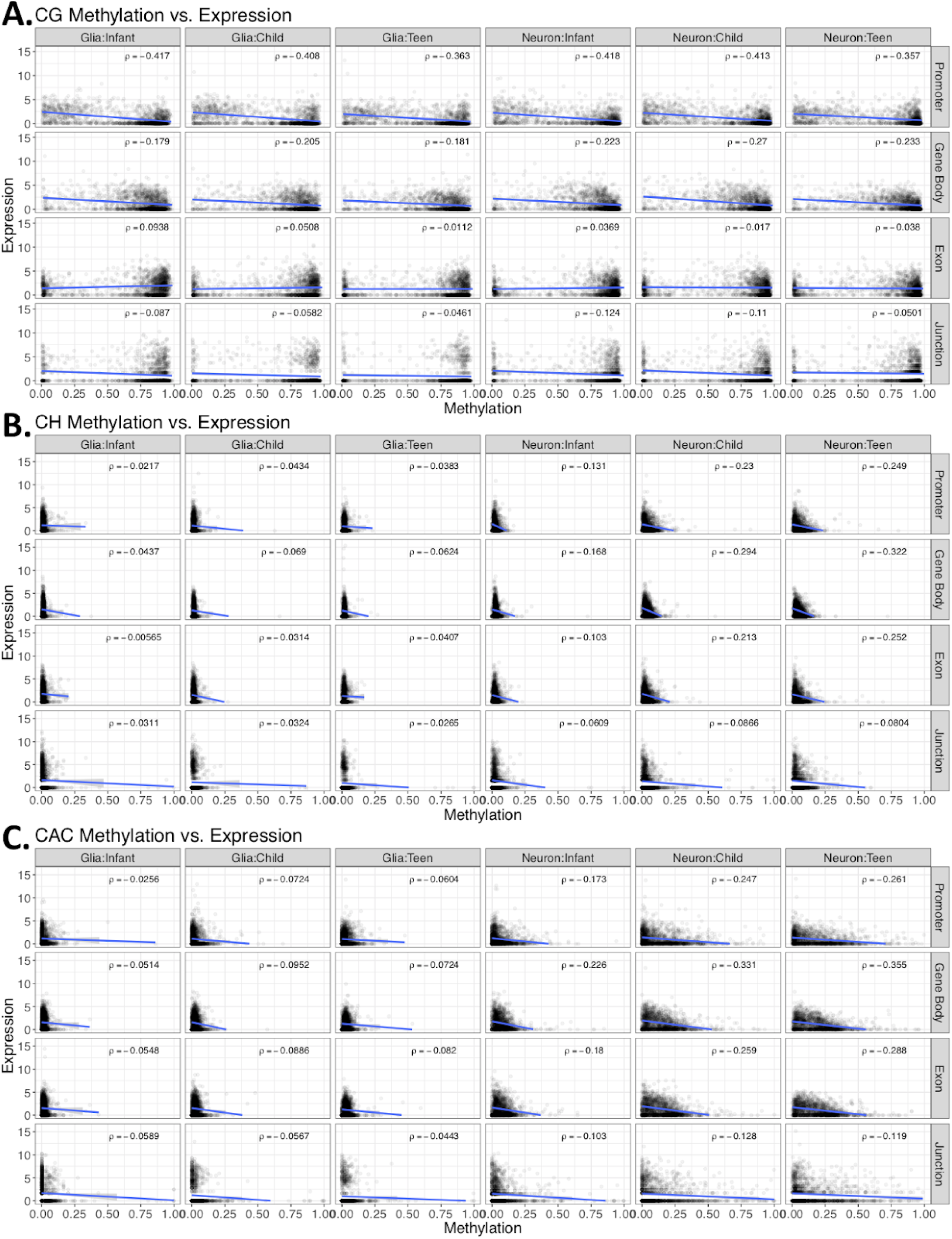
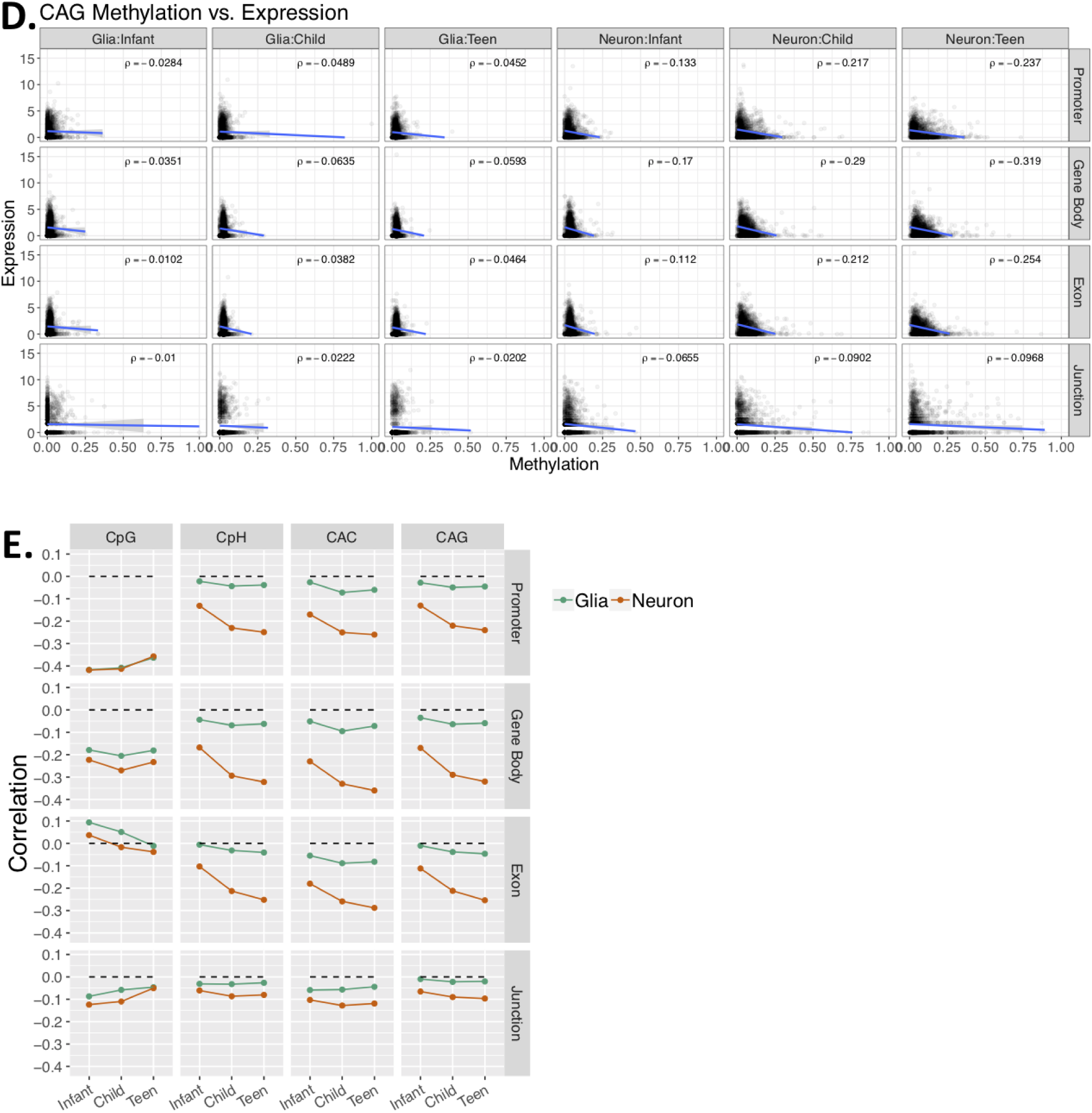
Relationship between methylation and expression. **(A-D)** A random sample of 10,000 expression feature:methylation pairs stratified by age and cell type in columns and feature type in rows. Each dot represents the mean methylation level of **(A)** CpGs, **(B)** CpHs, **(C)** CACs and **(D)** CAGs within the feature across all samples of that age and cell type on the x-axis and the log2([mean Rpkm]+1) for the promoters, gene bodies, and exons and the log2([mean junction-overlapping reads per 10 million mapped reads]+1) for the junctions on the y-axis. 10,000 pairs were chosen to reduce overplotting. Linear regression with shaded standard error and the rho for each correlation is listed for each plot. **(E)** Correlation of feature expression and methylation stratified by cytosine context (columns) and feature type (rows).

**Figure S13:**
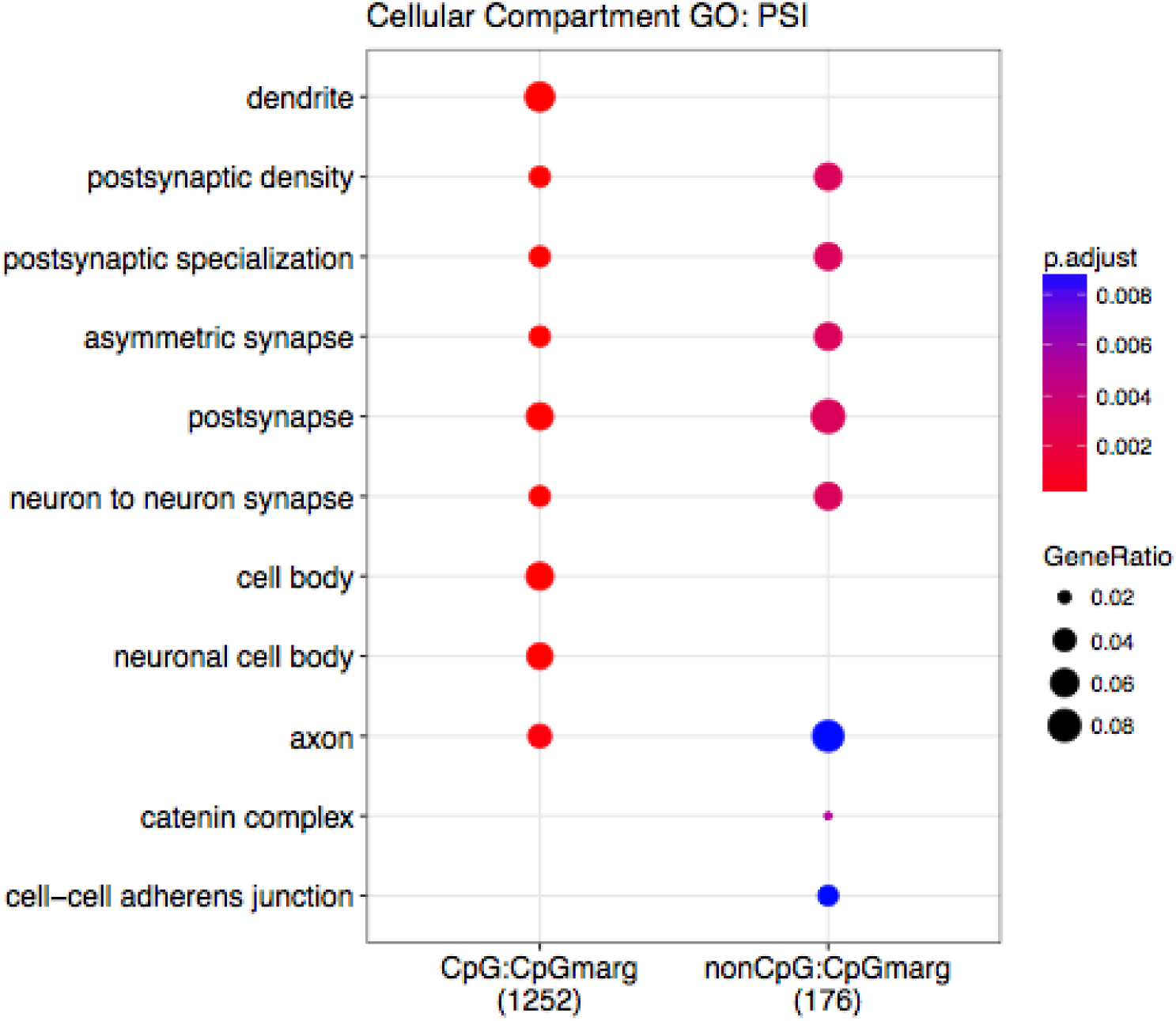
Cellular compartment ontology. Genes containing splicing events as measured by “percent spliced in” (PSI) that are associated with changing CpG and CpH methylation are enriched for cellular compartment gene ontology terms relating to neuronal features.

**Figure S14:**
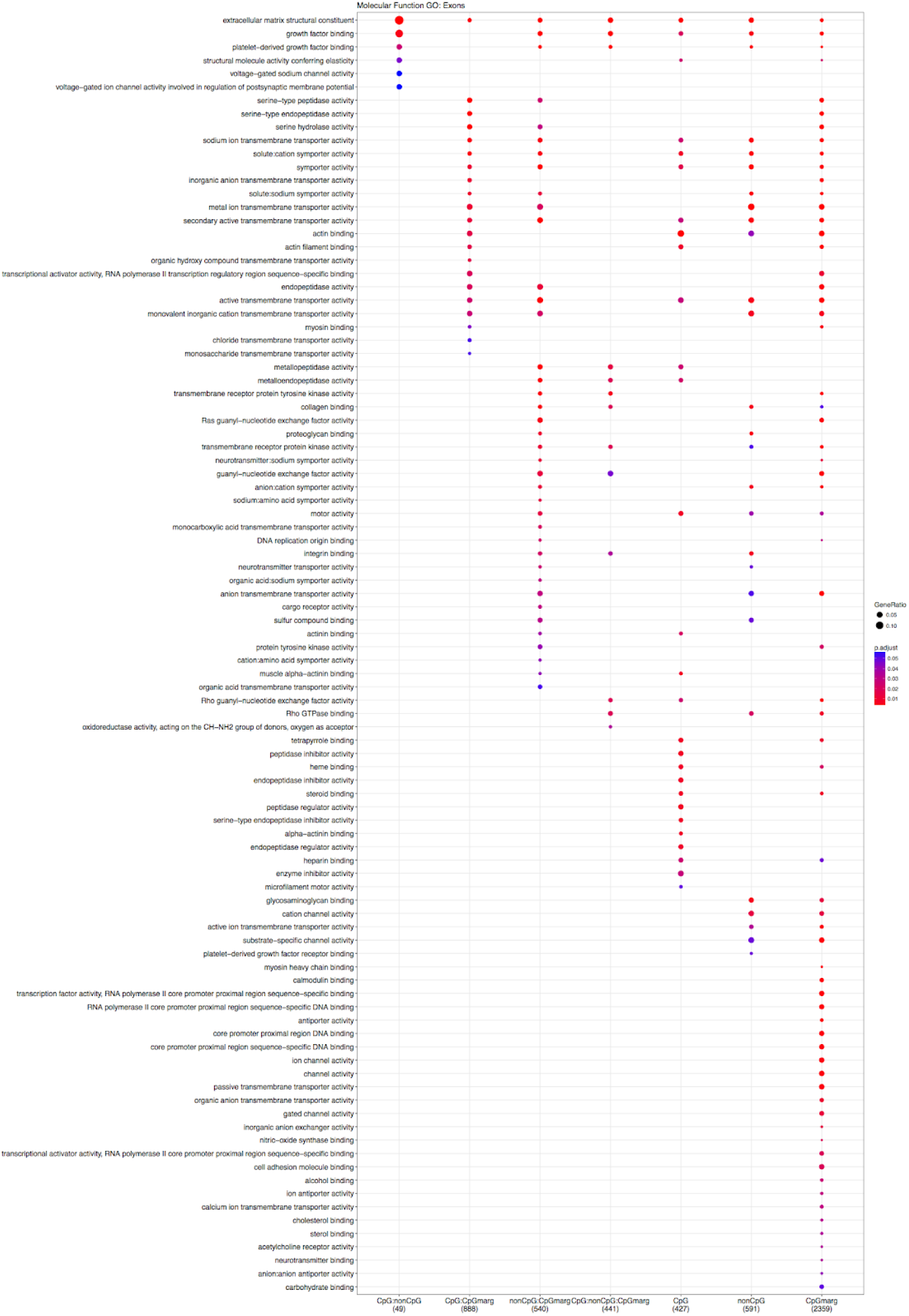
Molecular function ontology. Genes containing alternative exons that are associated with changing CpG and CpH methylation are enriched for molecular function ontology terms relating to neurons.

**Figure S15:**
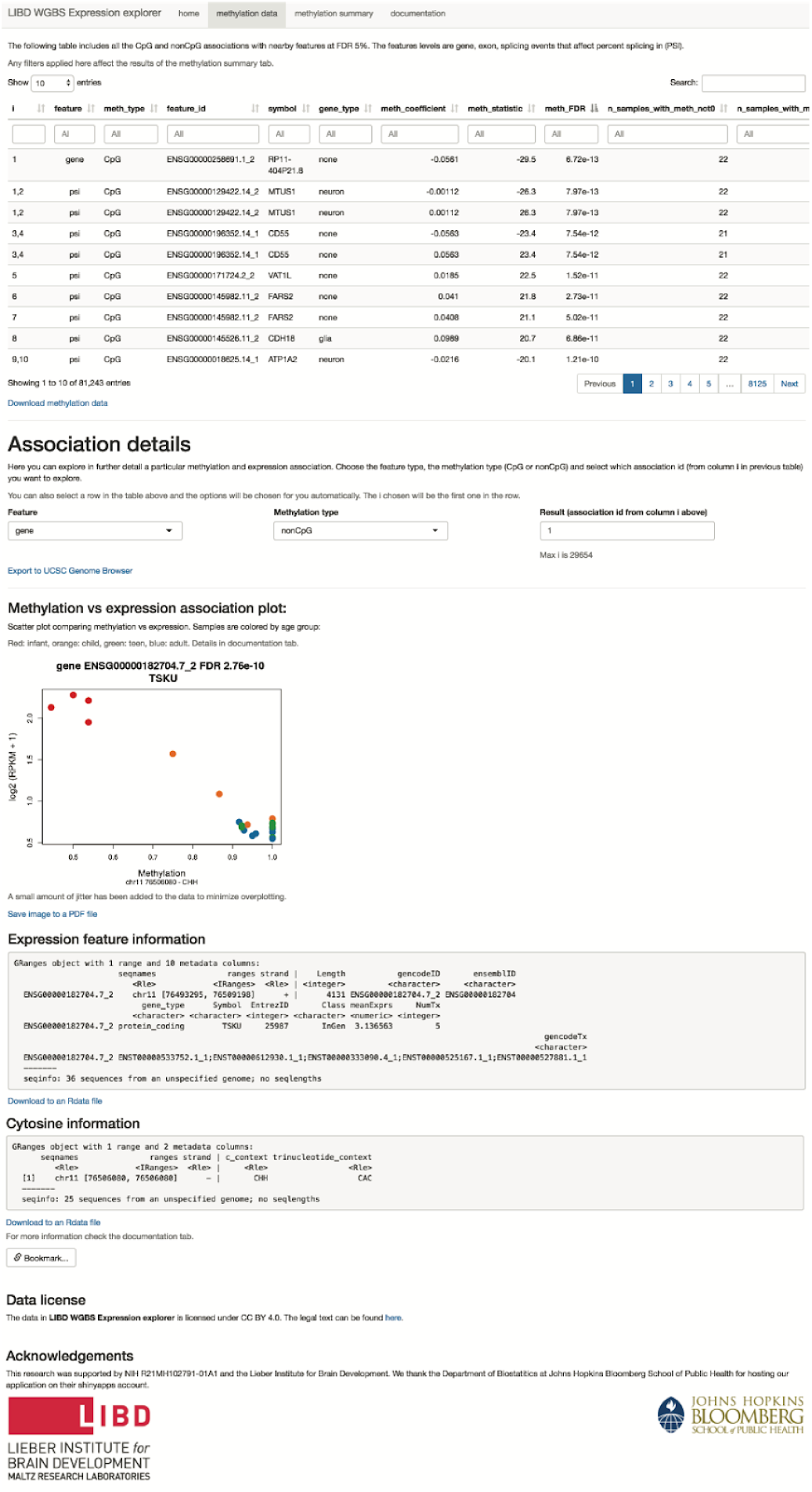
Web meQTL browser display. Interactive display of the CpHs and CpGs associated with expression at FDR<5% as shown at https://jhubiostatistics.shinyapps.io/wgbsExprs/. This screenshot shows the top nonCpG (mCpH) meQTL association at the gene expression level. Information about the gene is shown under *expression feature*, and information of the methylated C is shown under *cytosine*.

**Figure S16:**
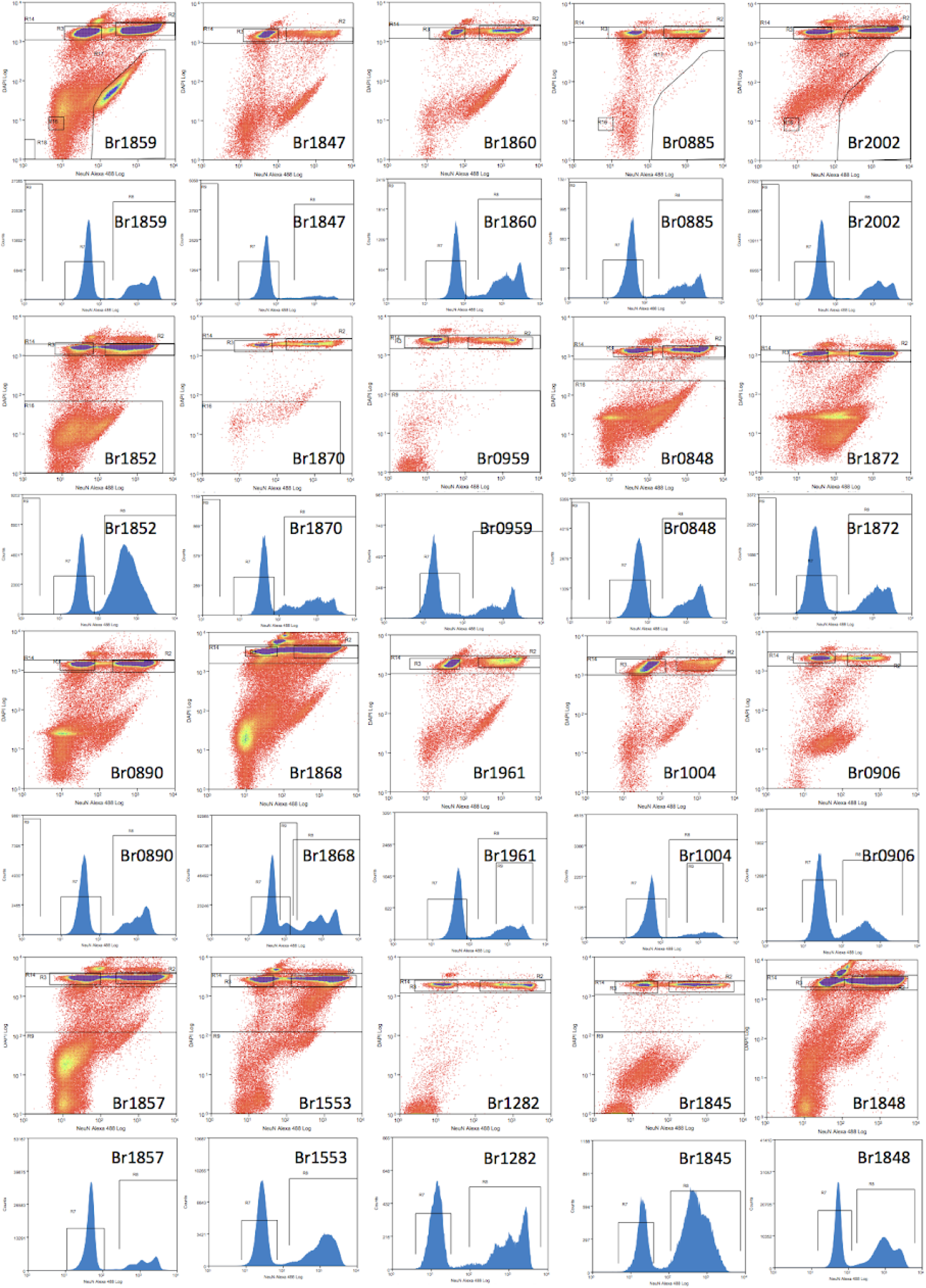
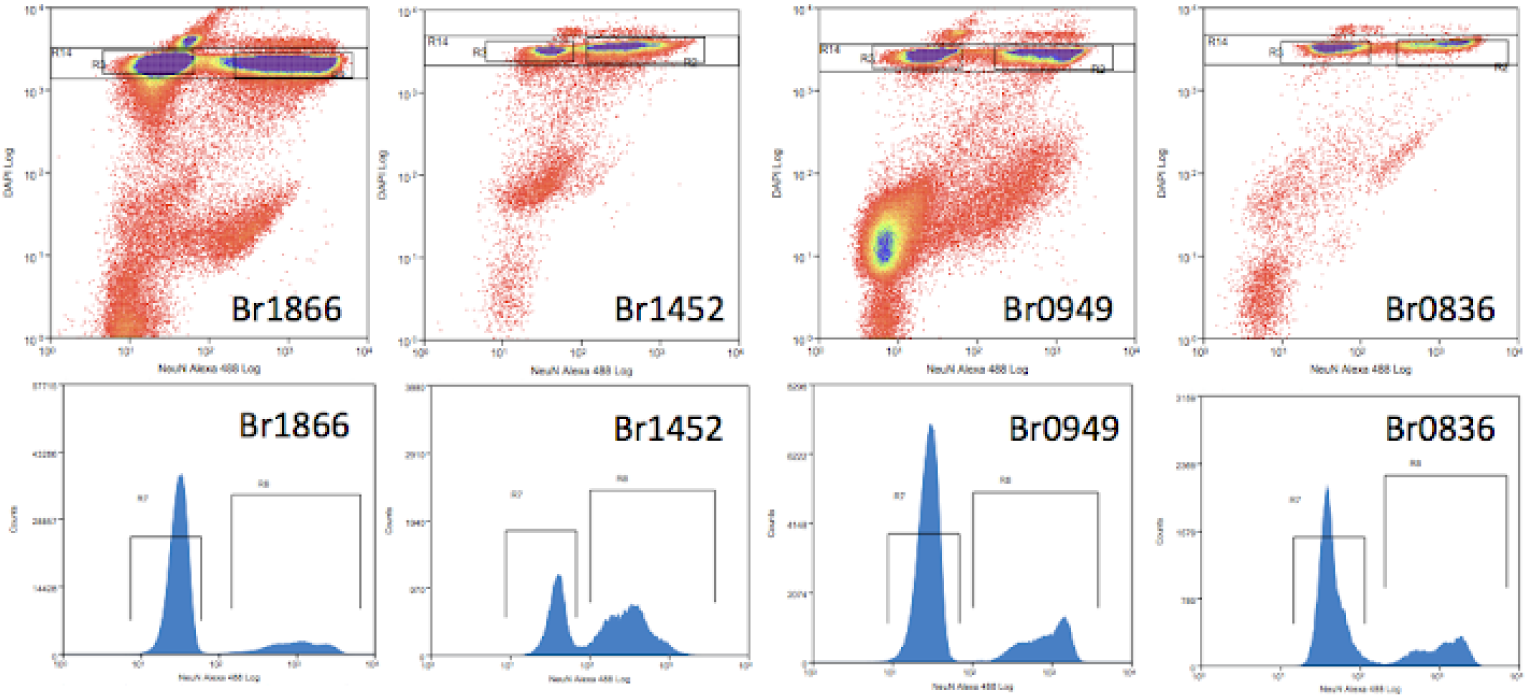
Raw sort data. Raw data collected from the MoFlo Legacy (Beckman Coulter) using Summit (version 4.3) software. The density plots show DAPI signal on the yaxis and Alexa Fluor 488-NeuN signal on the x-axis for events filtered via gates 1-3 as depicted in Figure S1A. The histograms show the distribution of Alexa Fluor 488-NeuN signal after including the gate for singlets based on DAPI signal (gate R14 in the raw data).

**Figure S17:**
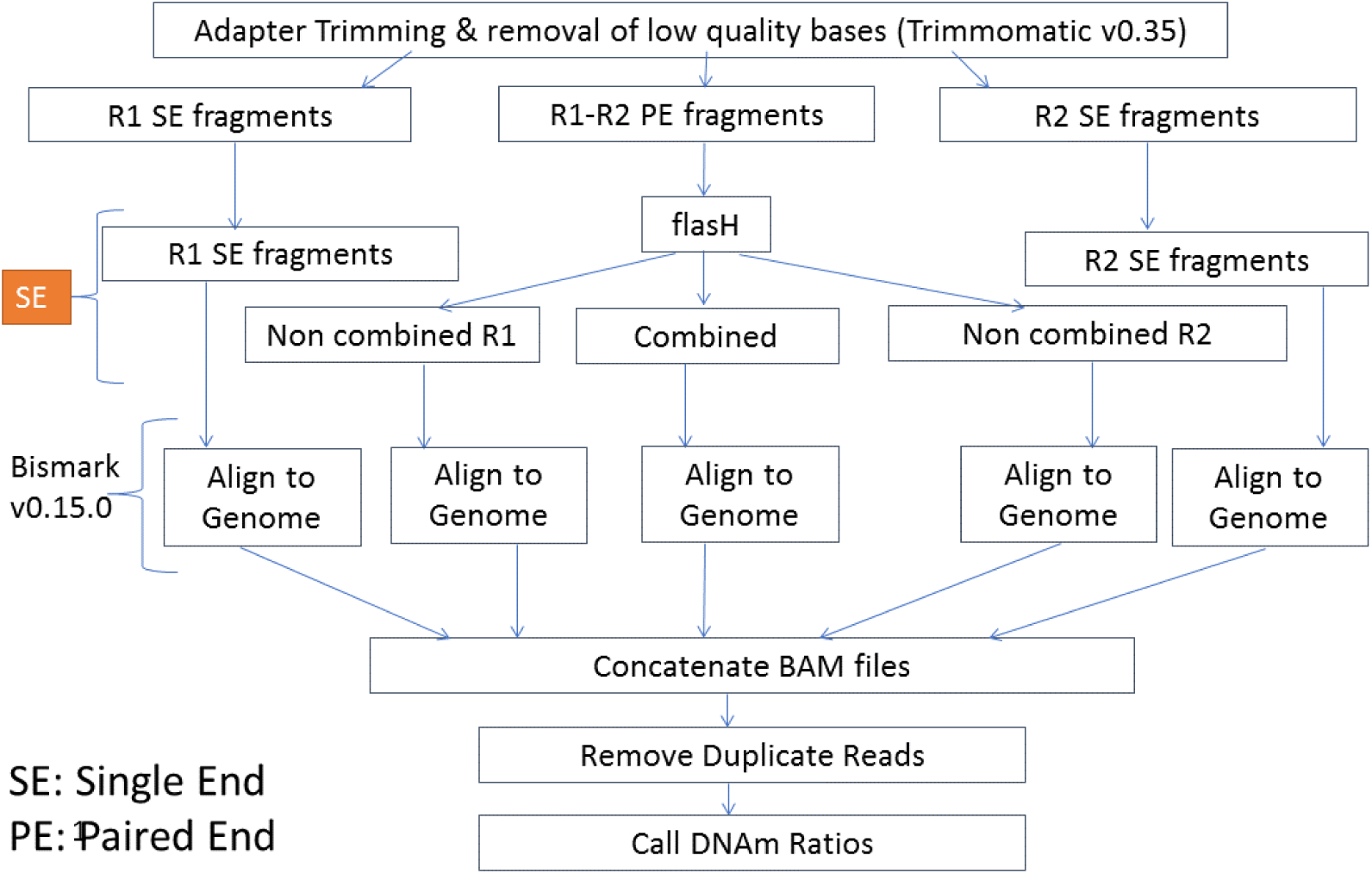
Data processing/alignment pipeline. Overview of the processing steps taken to prepare the WGBS data.

**Figure S18:**
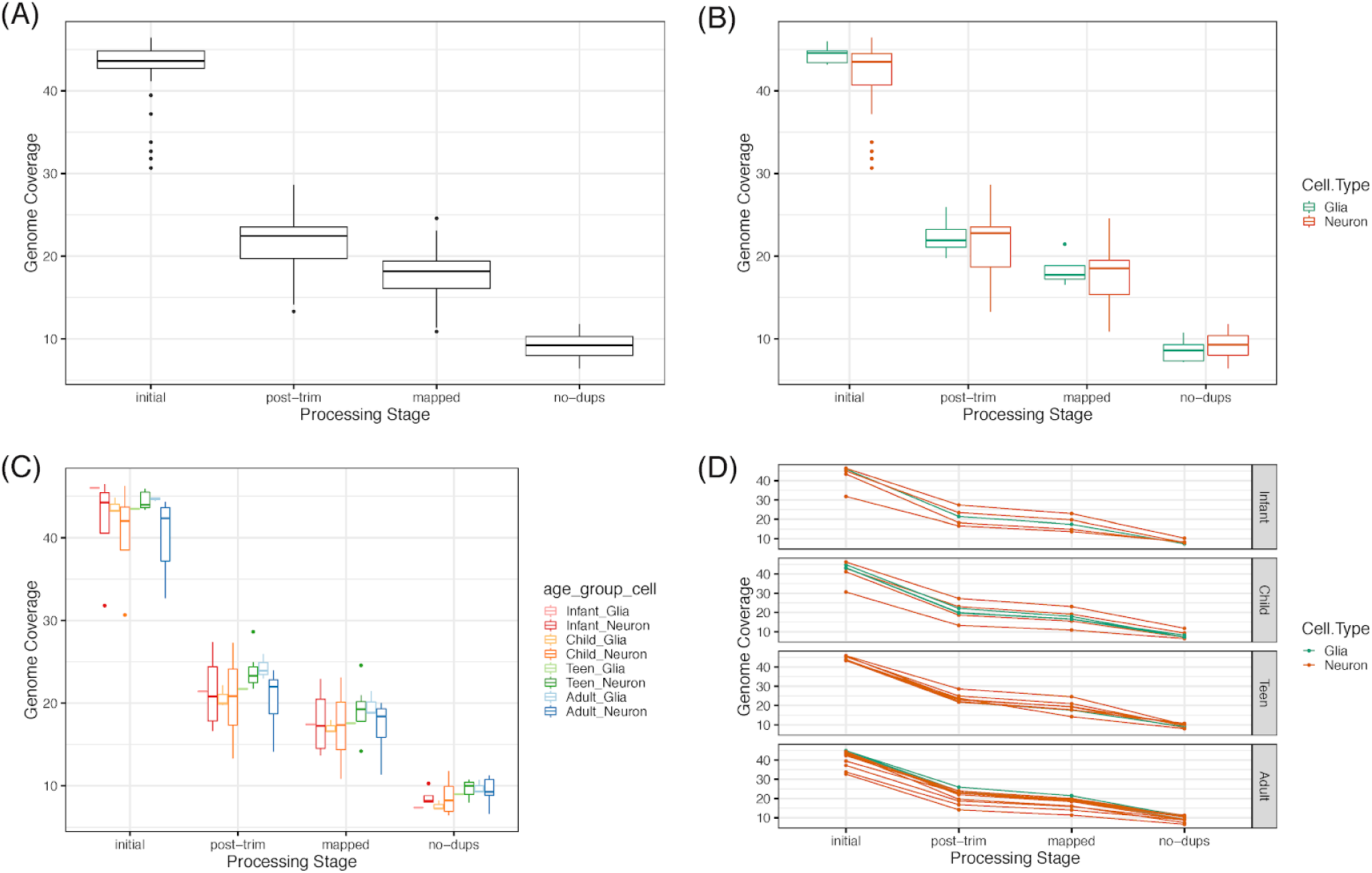
Genome coverage across processing stages. **(A)** Genome coverage across the four main processing stages: initial FASTQ files, after trimming, after mapping to the genome, and after removing duplicated reads. **(B)** Same as (A) but with samples separated by cell type. **(C)** Genome coverage across cell types and four age categories: infant, child, teen and adult. **(D)** Genome coverage trajectories for each sample separated by age category and colored by cell type.

**Figure S19:**
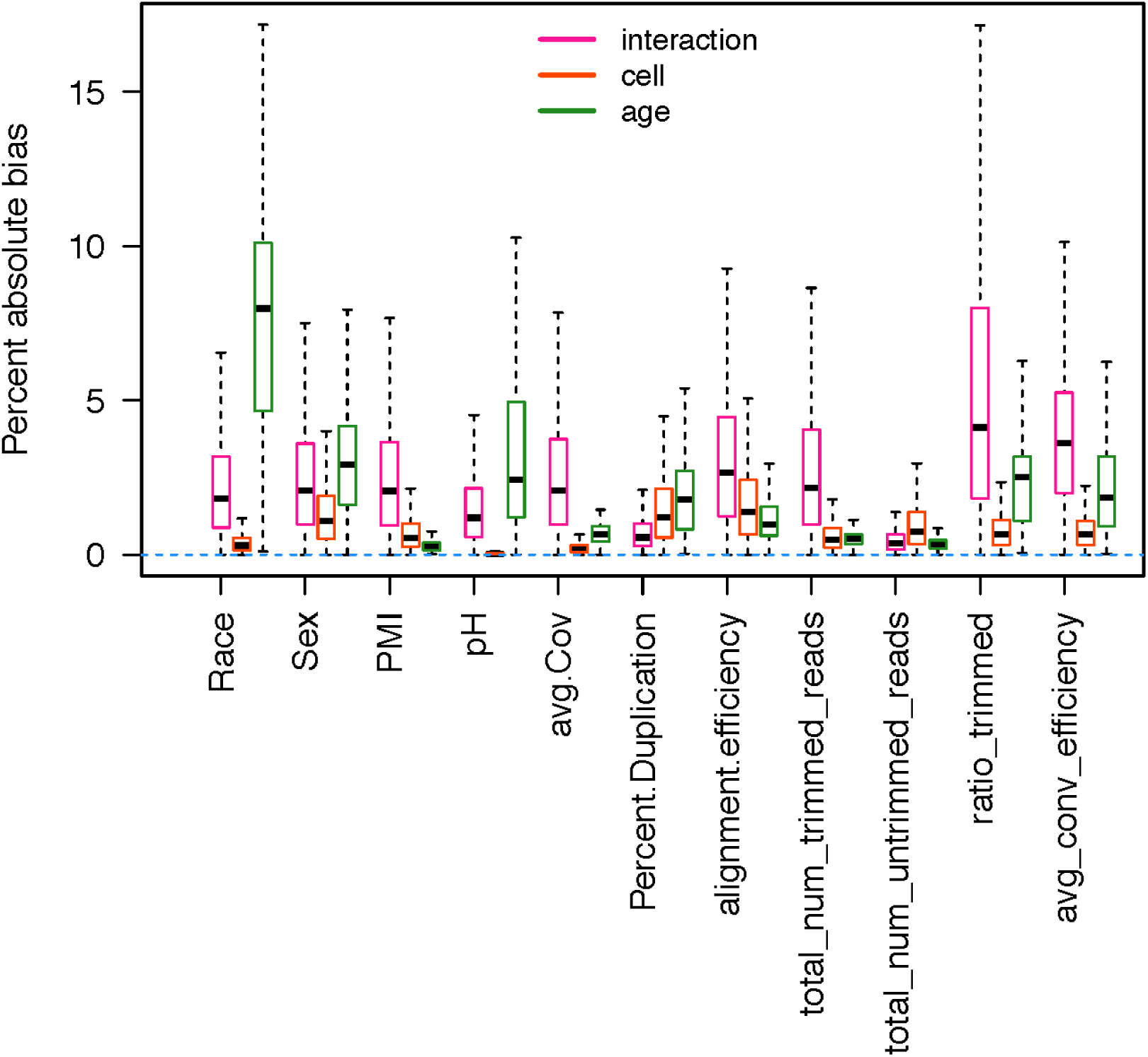
DMR sensitivity analyses. This figure shows the percent absolute bias for the three coefficients of interest used to define the DMRs: age (adjusting for cell type), cell type (adjusting for age), and the interaction between age and cell type. For each DMR type we computed the coefficients after adjusting for each of the covariates in the X-axis (*β*_adj_) and computed the percent absolute bias: 100 * |*β*_adj_ - *β*_original_ | / |*β*_original_|. All covariates considered show less than 10% absolute bias. The covariates considered are: race, sex, PMI, pH, average coverage, duplication percent, alignment efficiency, total number of trimmed reads, total number of untrimmed reads, ratio of trimmed reads, and average conversion efficiency (lambda).

**Figure S20:**
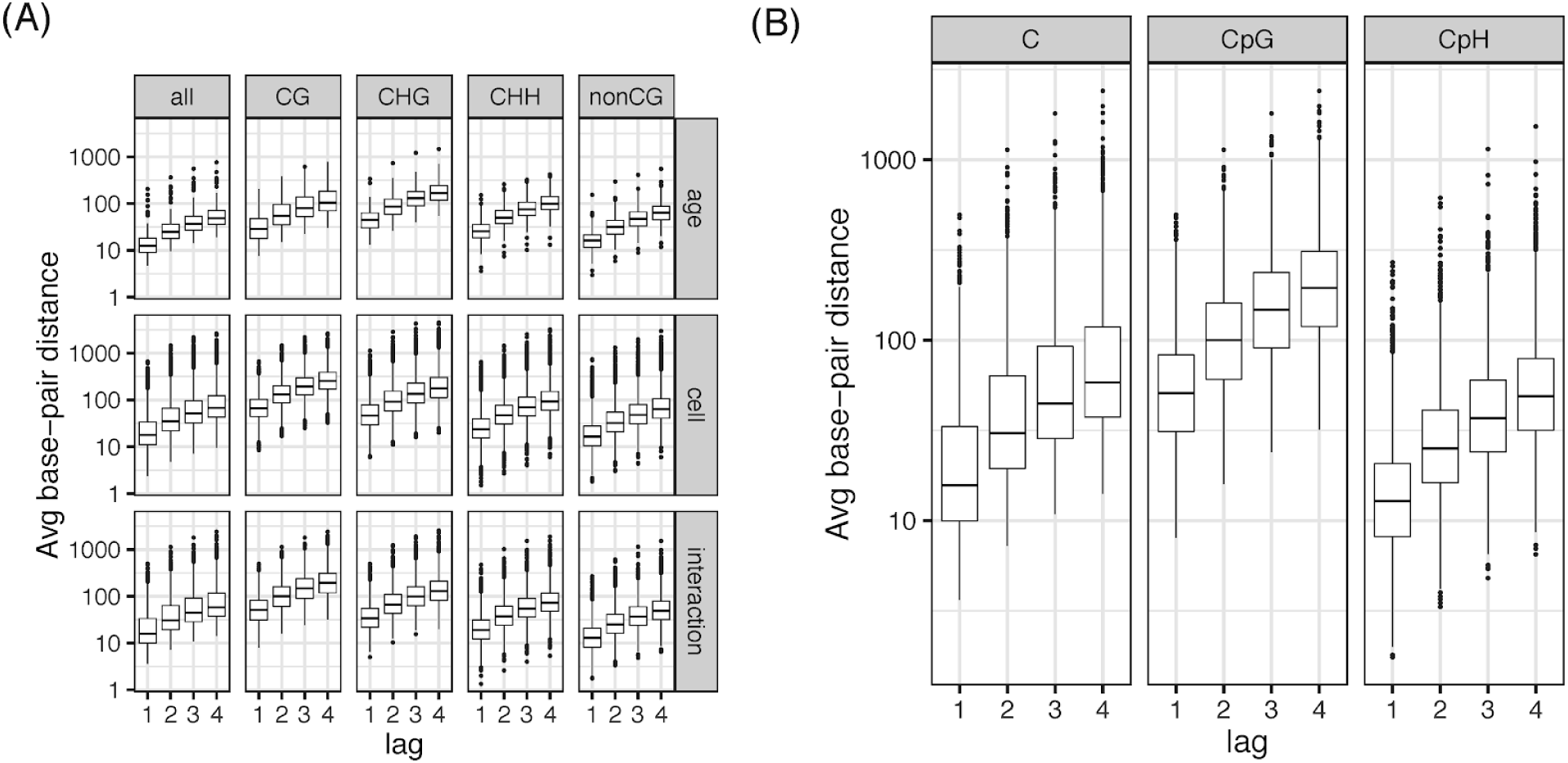
genome distance versus autocorrelation lag. For each of the cytosines considered for the autocorrelation analysis in Figure 2B, we computed the genome base-pair distance and averaged it for each group of cytosines. **(A)** Average genomic distance for each of the three DMR models and for various contexts of cytosines. **(B)** Average genome distance for the interaction DMRs (cdDMRs) for all cytosines, CpG and CpH. The genome distance is proportional to the lag. This panel is the exact complement to Figure 2B.

## Supplementary Tables

**Table S1: Phenotype and sequencing metrics for the WGBS samples.**

*See sheet 1 of Supplementary_tables.xlsx*.

**Table S2: Phenotype and sequencing metrics for the RNA-seq samples.**

*See sheet 2 of Supplementary_tables.xlsx*.

**Table S3:** Number of cytosines measured and distribution of methylation by cytosine context. Sample sizes for each sample type, number of CpGs and CpHs and distribution of CpG and CpHs across low, partial and high levels of methylation.

|  | N | CpG Context |  |  |  | CpH Context |  |  |  |
| --- | --- | --- | --- | --- | --- | --- | --- | --- | --- |
|  |  | # CpGs | Low (<20%) | Partial (20-80%) | High (>80%) | # CpHs | Low (<20%) | Partial (20-80%) | High (>80%) |
| Neuron | 24 | 18.7x10<sup>6</sup> | 10.10% | 13.90% | 76.00% | 58.1x10<sup>6</sup> | 91.60% | 8.00% | 0.40% |
| Glia | 8 | 18.7x10<sup>6</sup> | 11.10% | 17.80% | 71.10% | 58.1x10<sup>6</sup> | 99.40% | 0.50% | 0.10% |
| Fetal | 20 | 18.7x10<sup>6</sup> | 12.10% | 17.30% | 70.60% | 58.1x10<sup>6</sup> | 99.75% | 0.17% | 0.07% |
| Homogenate | 23 | 28.2x10<sup>6</sup> | 8.50% | 16.90% | 73.70% | 58.1x10<sup>6</sup> | NA | NA | NA |

**Table S4: Cell type-specific, developmental differentially methylated regions (cdDMRs).**

*See sheet 4 of Supplementary_tables.xlsx*.

**Table S5:** mC association with cell type and age in postnatal cell type-specific samples. Number of differentially methylated CpGs and CpHs across each of the statistical models: cell type effects adjusting for age, age effects adjust for cell type, cell type and age interaction effects, as well as age in neuron samples only.

| Model | CpG Context |  | CpH Context |  |
| --- | --- | --- | --- | --- |
| | FDR ≤ 0.05 | % Significant | FDR ≤ 0.05 | % Significant |
| **Cell type** | 4824804 | 25.80% | 7682075 | 18.80% |
| **Age** | 536164 | 2.90% | 3194618 | 7.80% |
| **Cell type:Age** | 90227 | 0.50% | 76 | 0% |
| **Age in Neurons** | NA | NA | 4020371 | 9.80% |

**Table S6: GWAS traits assessed using LDSC.**

*See sheet 6 of Supplementary_tables.xlsx*.

**Table S7: Stratified linkage disequilibrium score regression results.**

*See sheet 7 of Supplementary_tables.xlsx*.

**Table S8: Enrichment of DMRs and mCpH for disease-associated gene sets.**

*See sheet 8 of Supplementary_tables.xlsx*.

**Table S9: Enrichment for disease gene sets in DNAm-splicing association features.**

*See sheet 9 of Supplementary_tables.xlsx*.

**Table S10:** Variable dictionary for Table 1. Variables used in Table 1 with their full definition.

| Variable | Description |
| --- | --- |
| Feature | Either gene, exon or event affect the percent spliced in (PSI). |
| Methylation | Methylation type. Either CpG or CpH. |
| Direction | down. |
| N | Number of associations at FDR 5%. |
| N Unique C | Number of unique cytosines in these associations. |
| N Unique Features | Number of unique features in these associations. |
| N Meth>0 | Mean number of samples that have non-zero methylation values. Total 22. |
| N Meth<1 | Mean number of samples that have non-one methylation values. |
| Expr Change | Mean change in expression (ignoring NAs). |
| Age Confounded | Proportion of associations that are confounded by age, that is, proportion of associations where the methylation coefficient in a multiple linear regression adjusting for age has a FDR $\geq 5\%$. |
| Promoter | Proportion of cytosines in these associations that overlap gene promoters. A single cytosine can overlap multiple gene regions for different genes. |
| Gene Body | Proportion of cytosines in these associations that overlap gene bodies. |
| Gene Flanking | Proportion of cytosines in these associations that overlap gene flanking regions. |
| CHG | Proportion of cytosines in these associations that have a CHG context. |
| CHH | Proportion of cytosines in these associations that have a CHH context. |
| Glia | Proportion of features in these associations where the feature is differentially expressed between neurons and glia, with higher expression in glia. For genes and PSI, the gene list was used. Up to the top 5,000 features DE in each direction were used. |
| Neuron | Proportion of features in these associations where the feature is differentially expressed between neurons and glia, with higher expression in neurons. |
| No Diff | Proportion of features in outside the top differentially expressed genes between glia and neuron. |

